# Failed reprogramming of transformed cells due to induction of apoptosis and senescence impairs tumor progression in lung cancer

**DOI:** 10.1101/2023.10.19.563086

**Authors:** Pablo Pedrosa, Zhenguang Zhang, David Macias, Jianfeng Ge, Mary Denholm, Anna Dyas, Victor Nuñez-Quintela, Valentin Estevez-Souto, Patricia Lado-Fernandez, Patricia Gonzalez, Maria Gomez, Jose Ezequiel Martin, Sabela Da Silva-Alvarez, Manuel Collado, Daniel Muñoz-Espín

## Abstract

Cell reprogramming to pluripotency applied to the study of cancer has identified transformation and pluripotency as two independent and incompatible cell fates. A detailed knowledge of the relationship between transformation and reprogramming could lead to the identification of new vulnerabilities and therapeutic targets in cancer. Here, we explore this interplay and find that OSKM expression limits tumor cell growth by inducing apoptosis and senescence. We identify Oct4 and Klf4 as the main individual reprogramming factors responsible for this effect. Mechanistically, the induction of cell cycle inhibitor p21 downstream of the reprogramming factors acts as mediator of cell death and senescence. Using a variety of in vivo systems, including allografts, orthotopic transplantation and KRAS-driven lung cancer mouse models, we demonstrate that OSKM expression impairs tumor growth and reduces tumor burden.

## Introduction

The relation between pluripotency and cancer remains highly debated. The origin of the debate stems from the seminal works of Weinberg^1^ and Chang^2^ laboratories. In these two papers, the authors compared the transcriptional signatures of a variety of embryonic pluripotent stem cells, adult tissue-specific stem cells, and cancers of various degrees of malignancy. Their conclusion was that, although differing in many aspects, highly-malignant cells and embryonic pluripotent cells shared a common “ES-like signature”. These observations initially suggested that cancer cells achieve their maximal malignancy potential when they are less differentiated and resemble pluripotent states.

Nuclear reprogramming of differentiated cells to produce pluripotent embryonic stem cells (iPSCs or induced pluripotent stem cells) has been one of the greatest discoveries during the last decades^3^. This implies the ectopic expression of four essential factors, namely Oct4, Sox2, Klf4 and c-Myc (abbreviated here as OSKM), leading to a drastic change in cell identity. The application of cellular reprogramming to the study of cancer and to identify vulnerabilities in cancer cells is emerging as a potent strategy^4^. In particular, an interesting aspect is the similarity between reprogramming to pluripotency and neoplastic transformation^5^. Remarkably, reprogramming factors can act in cancer as oncogenes. For instance, Oct4 drives the transformation potential of testicular germ cells^6^; Sox2 is an amplified lineage-survival oncogene in lung and oesophageal squamous cell carcinomas^7,8^; Klf4 can function as a dominant oncogene, as it is frequently overexpressed in human breast tumors and squamous cell carcinomas^9^; and c-Myc is a well-known gene deregulated in multiple human cancers^10^. Both processes, reprogramming to pluripotency and neoplastic transformation, involve the acquisition of new epigenetic programs^11^. Accordingly, several chromatin modifiers acting on reprogramming have also been found operating in cancer^12^.

The similarities can also be extended to common barriers featured by the processes of reprogramming and transformation. For both, the activities of several tumor suppressor genes such as *Ink4/Arf* and *p53* result in decreased generation of pluripotent cells by normal differentiated cells and reduced capacity of transformation^13–18^. One key process that contributes to overcome the barriers that impair the transformation into a cancer cell is an initial step of immortalization. It would therefore be expected that transformed cells are more susceptible to being reprogrammed. However, there are very few reported examples of tumor cells capable of undergoing full stable reprogramming into pluripotency, mainly in hematopoietic cancer cells with single or few oncogenic events driving tumorigenesis^19,20^. Some other cases of partial or limited reprogramming include melanoma^21^, gastrointestinal cancer cells^22^ and pancreatic ductal adenocarcinoma cells^23^. In most of the cases though, cancer cells remain resistant to being reprogrammed^20^. A recent work by Ito et al.^24^ emphasizes the oncogene-dependent resistance to reprogramming and how the study of the underlying mechanisms can be exploited to identify therapeutic targets.

In line with these results, we previously tested the consequences of introducing an activated RAS oncogene as part of the reprogramming cocktail of Oct4, Sox2, Klf4 and c-Myc^25^. Interestingly, we found that while RAS increases the reprogramming efficiency in normal cells its expression in the context of cellular transformation impairs reprogramming. However, when oncogenic RAS is switched off the capacity of reprogramming is fully restored, suggesting that cancer and reprogramming are incompatible cellular fates. Supporting this concept others have also recently found that the molecular trajectories of reprogramming and oncogenic transformation, although initially shared, diverge into two separate and defined cell entities^26^.

Here, we have further explored this relationship between oncogenic transformation and cell reprogramming by analyzing the effect of expressing the reprogramming factors on lung cancer cells. Our experiments confirm that both human and mouse lung cancer cells are highly resistant to reprogramming. We find that OSKM expression limits tumor cell growth by inducing apoptosis and senescence, and dissect the specific contribution of the independent factors. Mechanistically, we identify the induction of the cell cycle inhibitor p21 downstream of the reprogramming factors as a mediator of this effect. We further validate our findings in vivo using allograft and orthotopic transplantation of lung tumor cells as well as a genetically modified mouse model of KRAS-driven lung cancer. Our data show that OSKM expression in lung cancer cells does not lead to pluripotency, but rather affects tumor growth and reduces tumor burden.

## Results

### OSKM expression impairs the growth of lung cancer cells

Immortalization removes a barrier for cellular reprogramming by OSKM allowing a more efficient de-differentiation and iPSC generation^13–18^. However, complete neoplastic transformation, which requires previous immortalization, impairs successful reprogramming, suggesting that reprogramming and transformation are incompatible cell fate^25^. We further explored this by subjecting different transformed cells to the expression of OSKM and confirmed the inefficient reprogramming of these cells (**Supplementary Fig. 1**). During these experiments we typically observed that cell cultures expressing OSKM were negatively affected by the expression of the reprogramming factors. This prompted us to analyze in more detail the effect of expressing the reprogramming factors in cancer cells.

We focused on lung cancer cells A549 and L1475luc (derived from a mouse model expressing oncogenic Kras and lacking p53, namely KP) cells^27^ of human and mouse origin, respectively. Both cell lines were transduced with lentiviruses or transposon vectors that express OSKM on a doxycycline-inducible manner (**Fig. 1a**). We confirmed the upregulation of the reprogramming factors by RT-qPCR and Western blot in A549 cells (hereafter named A549-rtTA-OSKM) (**Supplementary Fig. 2a**). For the L1475luc cells (hereafter named L1475luc-rtTA-OSKM), after transfection mOrange positive cells (expressing OSKM factors) were sorted by flow cytometry and the expression of Oct4 was confirmed by RT-qPCR and Western blot (**Supplementary Fig. 2b, c**). When doxycycline was added for three days to both cell lines, we observed a reduced growth of the cell cultures (**Fig. 1b**). Analysis at longer times by colony formation assay confirmed the impaired growth caused by OSKM expression (**Fig. 1c**). A549-rtTA-OSKM cells also showed a reduced capacity to form colonies in soft agar when expressing OSKM (**Supplementary Fig. 2d**), and when they were allowed to form colonies, addition of doxycycline to induce the expression of OSKM caused their destruction (**Fig. 1d**). We added to these colonies a probe that allows the detection of the active form of Caspase-3, indicative of apoptosis induction. Expression of the reprogramming factors increased the fluorescent signal indicating an increase in apoptosis coinciding with the destruction of the colonies (**Fig. 1d**).

**Fig. 1:**
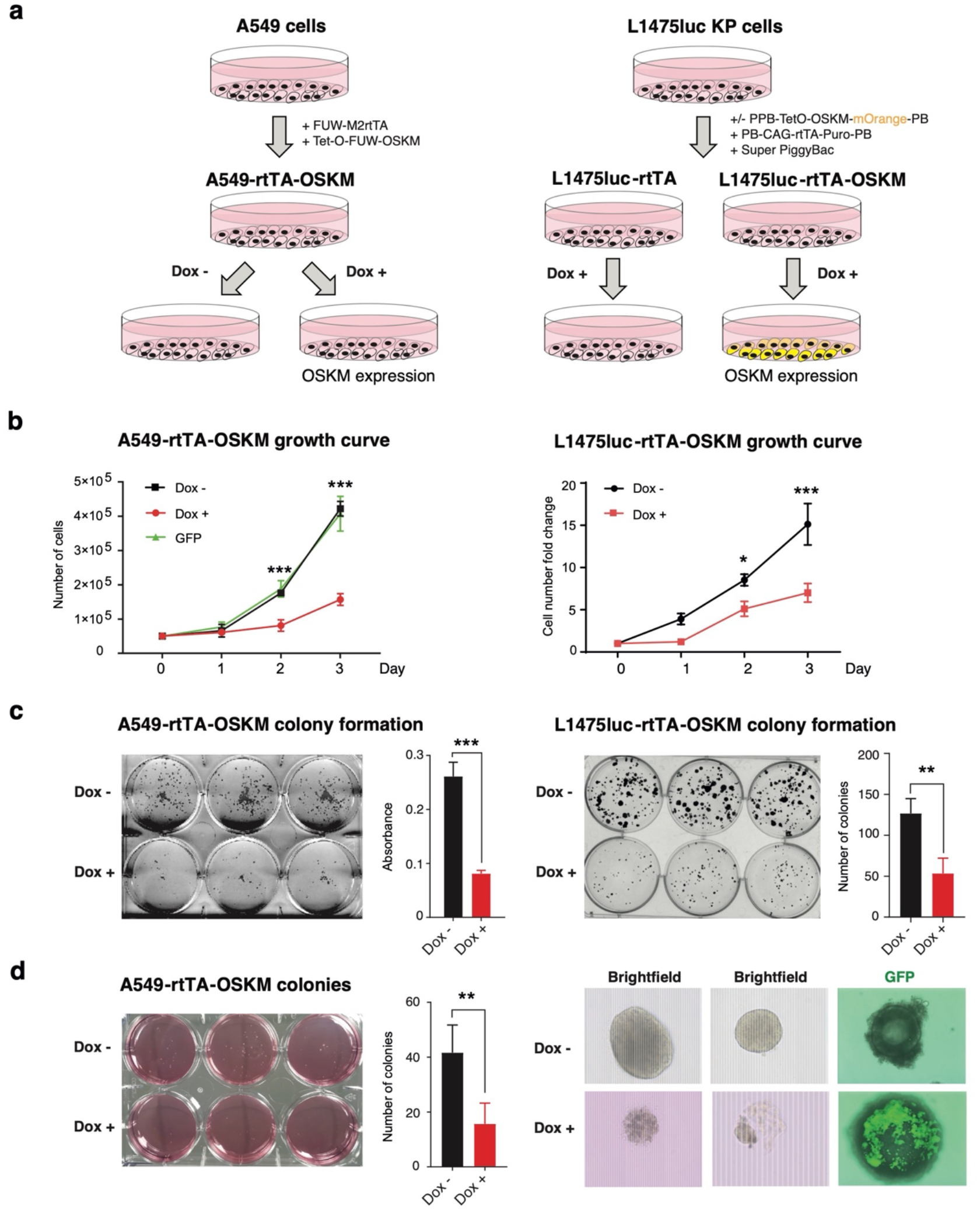
OSKM expression reduces proliferation and colony formation in lung adenocarcinoma cell lines A549 and L1475luc. **(a)** Schematic representation of the generation of the experimental systems. Left: Lentiviral vectors carrying the reprogramming factors (OSKM) and the tetracycline-dependent reverse transactivator (rtTA) were transduced into A549 cells, allowing the expression of the reprogramming factors in the presence of doxycycline in the culture medium. The conditions to be analyzed are therefore Dox + (expression of the factors) and Dox - (no expression of the factors). Right: For OSKM expression in L1475luc cells, three transposon vectors containing TetO-OSKM-mOrange, rtTA and PiggyBac were transfected into the cells. For single rtTA expressing cells, only the latter two vectors were used. **(b)** Cell growth curves of A549-GFP and A549-rtTA-OSKM cells (left) or L1475luc rtTA-OSKM cells (right), over 3 days, treated or not with doxycycline (1 µg/ml). **(c)** Representative images and quantifications of colony formation assays of A549-rtTA-OSKM cells (left) or L1475luc-rtTA and L1475luc-rtTA-OSKM cells (right), over 3 days, treated or not with doxycycline (1µg/ml). Mean and SD are shown. **(d)** Left: Representative image and quantification of a colony formation assay in soft agar of A549-rtTA-OSKM cells treated or not with doxycycline (1µg/ml). Right: Representative images of colonies of A549-rtTA-OSKM cells treated or not with doxycycline (1µg/ml), and detection of cleaved Caspase-3 using a fluorescent probe. Statistical significance was calculated using Student’s t-test, ***P<0.001; **P<0.01; *P<0.05. Data are mean ± SD.

Altogether, these data show that OSKM expression results in a reduced growth of both human and mouse lung cancer cells.

### Expression of the reprogramming factors activate apoptosis and cellular senescence in lung cancer cells

Next, we focused on analyzing the impact of expressing the reprogramming factors on the lung cancer cell fate. Since we observed an increase of a fluorescent probe for the detection of active Caspase-3, suggesting the induction of apoptosis, we decided to analyze this cell response by flow cytometry using antibodies against cleaved Caspase-3 and for Annexin V. A549 cells expressing the reprogramming factors showed an increase in both apoptotic markers compared with the control condition (**Fig. 2a**). To further confirm the involvement of apoptosis in the observed impaired growth, we treated cells expressing OSKM with a pan-Caspase inhibitor to block the induction of apoptosis. First, we confirmed that incubation with the Caspase inhibitor resulted in a reduced activation of cleaved Caspase-3 (**Supplementary Fig. 3a**). Then, we followed cell cultures expressing the reprogramming factors in the presence or absence of this inhibitor and observed a partial recovery of cell growth at short (3 days) and long time points (14 days) when the pan-Caspase inhibitor was added to the OSKM-expressing cell cultures (**Fig. 2b**). Similarly, when mouse cancer cells incubated with an apoptosis fluorescent reporter were followed using a live cell analysis system, we confirmed the impaired growth and apoptosis induction caused by OSKM expression (**Fig. 2c**). This reduced growth does not reflect an intrinsic different proliferative capacity of both cell lines since they showed a similar growth rate in the absence of doxycycline (**Supplementary Fig. 3b**). The reduction in cell proliferation and the induction of apoptosis were further confirmed by flow cytometry (**Supplementary Fig. 3c, d**).

**Figure 2.**
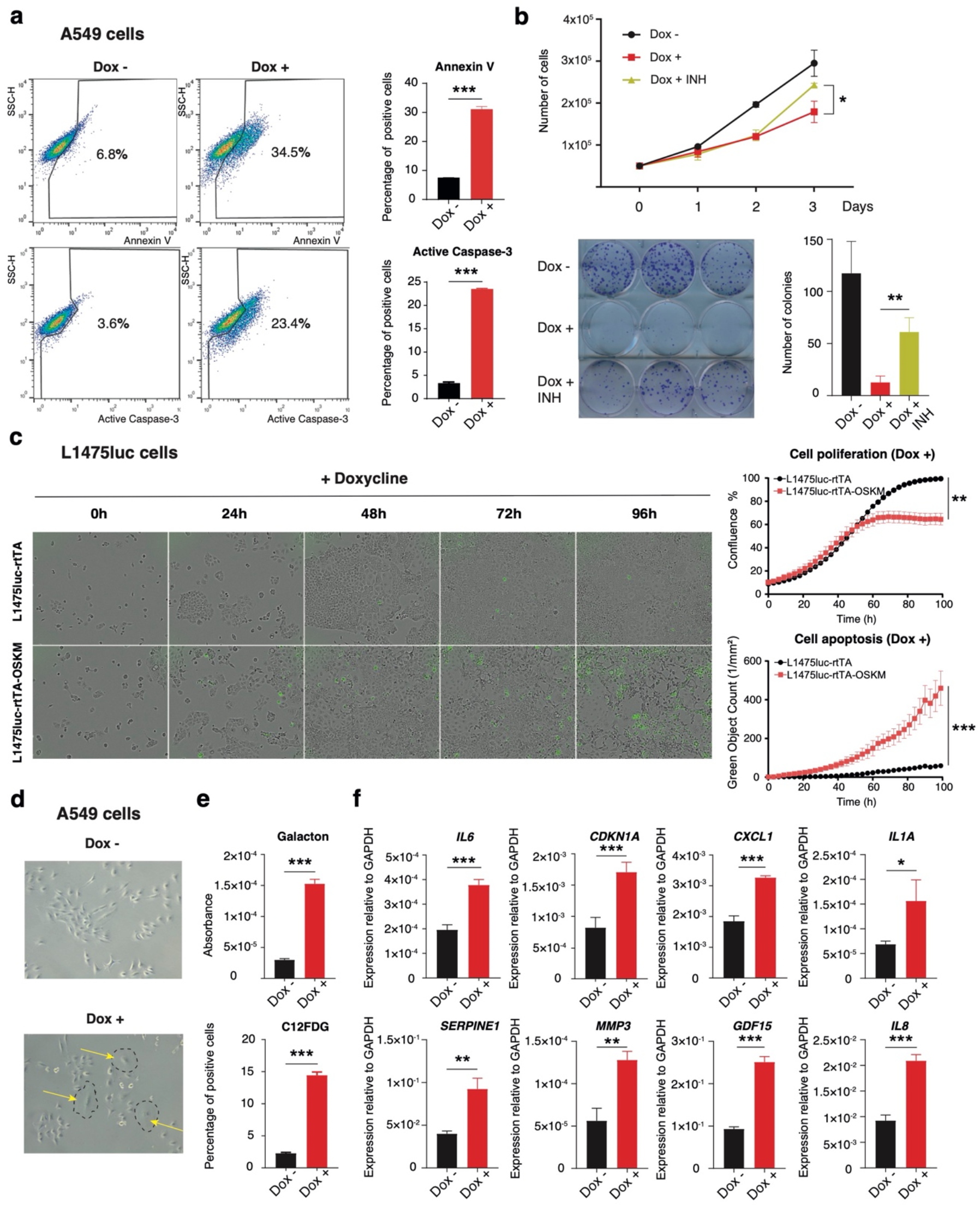
OSKM expression induces lung cancer cell apoptosis and senescence in vitro. **(a)** Flow cytometric analysis of Annexin V levels (left top) and quantification (right top) and of cleaved Caspase-3 levels (left bottom) and quantification (right bottom). **(b)** Analysis of the proliferative capacity (top) and clonal expansion (bottom) in A549-rtTA-OSKM cells expressing (Dox +) or not (Dox -) the reprogramming factors, and treated or not with the caspase inhibitor (INH). **(c)** Quantification (top panels) and representative images (bottom panels) of cell proliferation and apoptosis in Incucyte experiments. Cells were treated or not with 1 µg/ml doxycycline and images taken every 3h for over 96h. Incucyte Caspase-3/7 Green Dye was used to detect green apoptotic cells. Data are mean ± SD (n=3). **(d)** Representative microscopy images of A549-rtTA-OSKM cells expressing (Dox +) or not (Dox -) the reprogramming factors. Arrows in yellow point to cells showing typical enlarged and flattened morphology of senescent cells (indicated by dashed lines). **(e)** Measurement of SA-ß-gal activity using Galacton adjusted to the number of cells (top) or C12FDG by flow cytometry (bottom). **(f)** Expression levels by RT-qPCR of mRNAs for: *IL6*, *CDKN1A*, *CXCL1*, *IL1A*, *SERPINE1*, *MMP3*, *GDF15* and *IL8*. Statistical significance was calculated using Student’s t-test, ***P<0.001; **P<0.01; *P<0.05. Data are mean ± SD.

While we were performing these analyses, we observed the frequent appearance of cells showing the typical cell senescence morphology in the cell cultures of A549 cells expressing OSKM. Cells were enlarged and flattened, and showed multivesicular cytoplasms (**Fig. 2d**). To confirm and quantify this observation, we performed two assays to measure the levels of senescence-associated beta-galactosidase activity (SA-ß-gal), the most widely used marker of cell senescence. Using two different substrates: Galacton, a chemiluminescent substrate; and C12FDG, a fluorescent one, we confirmed the induction of cell senescence after expression of the reprogramming factors (**Fig. 2e**). In addition, we measured the mRNA levels of several markers of cell senescence (*IL6*, *CDKN1A*, *CXCL1*, *IL1A*, *SERPINE1*, *MMP3*, *GDF15* and *IL8*) and observed an increased expression after OSKM induction (**Fig. 2f**).

The above results indicate that the expression of the reprogramming factors on lung cancer cells triggers cell defense responses such as apoptosis and senescence that impair cell growth.

### Transcriptomic and proteomic profiles reinforce the induction of apoptosis and senescence upon OSKM expression

To gain mechanistic insights on how gene expression was affected by OSKM induction, we performed bulk RNA-seq transcriptomic analyses utilizing both human A549-rtTA-OSKM and mouse L1475luc-rtTA-OSKM lung cancer cell lines and their control counterparts (lacking the OSKM cassette) upon doxycycline exposure. Initial validation by RT-qPCR confirmed the increased expression levels of mRNA corresponding to the four Yamanaka factors in A549-rtTA-OSKM cells (**Supplementary Fig. 2a**) and L1475luc-rtTA-OSKM cells (**Supplementary Fig. 4a**). Accordingly, gene set enrichment analysis (GSEA) of the transcriptomic data showed that the Hallmark of MYC targets V1 and V2 are both significantly upregulated in human A549-rtTA-OSKM and mouse L1475luc-rtTA-OSKM lung cancer cell lines (**Fig. 3a, b** and **Supplementary Fig. 4b, c**), consistent with OSKM expression. KEGG pathway database analyses showed that signalling pathways regulating Pluripotency of Stem Cells and Apoptosis are significantly upregulated in both human and mouse lung cancer cell lines suggesting that the induction of pluripotency features by OSKM overexpression and the activation of apoptosis could be functionally interconnected (**Supplementary Table 1**). In addition to apoptosis, pathways of cellular senescence were also significantly upregulated in mouse L1475luc-rtTA-OSKM lung cancer cells when compared to their control counterparts (**Supplementary Table 1**). Next, we reanalyzed the significantly upregulated genes from the Apoptosis and Signalling Pathways Regulating Pluripotency of Stem Cells in KEGG using a different analysis platform, namely WikiPathways^28^. Enrichment plots (Emapplot and Circular cnetplot) showed that pathways of Apoptosis and ESC Pluripotency Pathways are significantly upregulated in A549-rtTA-OSKM *vs* A549-rtTA cells (**Fig. 3a**) and L1475luc-rtTA-OSKM *vs* L1475luc-rtTA cells (**Fig. 3b**) upon doxycycline exposure. Our analyses showed a significant enrichment of genes encoding for known apoptosis mediators, such as *FAS* and *CASP9* genes in human A549-rtTA-OSKM cells or *Bid* in mouse L1475luc-rtTA-OSKM lung cancer cells (**Fig. 3a, b**). Furthermore, upregulation of pathways (such as p53 signalling) and enrichment of genes, such as *Cdkn1a* (encoding p21), related to the implementation of senescence, or *IL6* or *IL1A,* related to the senescence-associated secretory phenotype (SASP), were observed in L1475luc-rtTA-OSKM cells *vs* L1475luc-rtTA cells or A549-rtTA-OSKM *vs* A549-rtTA cells, respectively, upon doxycycline addition (**Fig. 3a, b**). This correlates with the significant upregulation of p53 pathway, NFkB, and MTOR signaling hallmarks, all of them often activated in senescent cells, detected in the GSEA analyses (**Supplementary Fig. 4b, c**).

**Figure 3.**
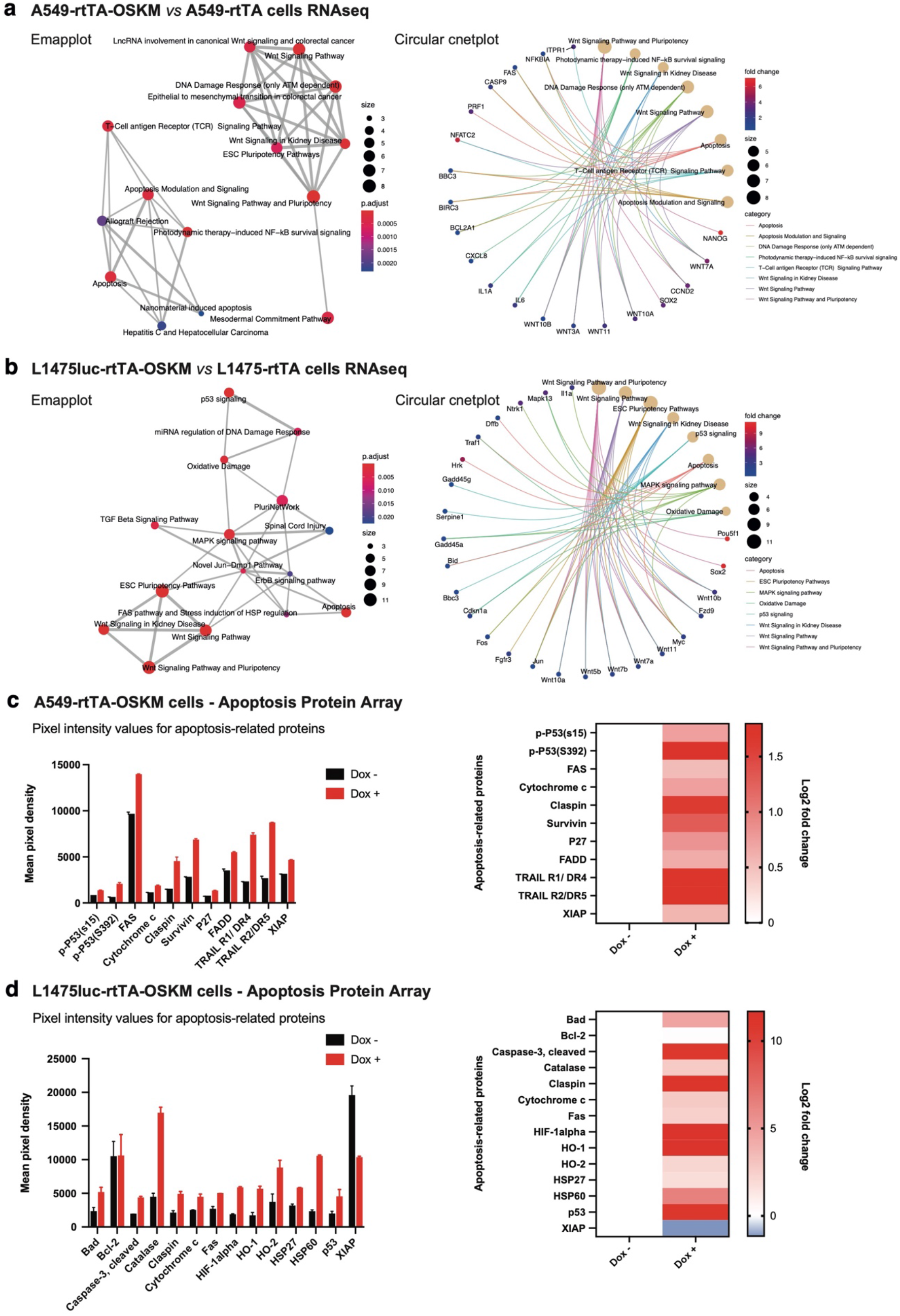
Transcriptomic and proteomic profiles showing evidence of apoptosis and senescence induction in lung cancer cells upon OSKM expression. Enrichment plots (Emapplot, left and Circular cnetplot, right) showing upregulated pathways and genes in A549-rtTA-OSKM *vs* A549-rtTA cells **(a)** and L1475luc-rtTA-OSKM *vs* L1475luc-rtTA cells **(b)** upon doxycycline exposure. Emapplot of enriched “Biological Process” gene ontology terms (P < 0.05, FDR < 0.05). p.adjust = the Benjamini-Hochberg adjusted P-value for the enriched ontology term. Fold change = the fold change difference in the annotated genes between OSKM-expressing cells and control cells. Size = the number of differentially expressed genes which belong to the enriched gene ontology term or category. **(c)** Graph showing pixel intensity values using a human apoptosis-related protein array with A549-rtTA-OSKM cells expressing (Dox +) or not (Dox -) the reprogramming factors (left) and associated heatmap (right). **(d)** Same as (c) but using a specific mouse apoptosis protein array for L1475luc-rtTA-OSKM cells expressing (Dox +) or not (Dox -) the reprogramming factors. Log2 fold change = the logarithm to base 2 fold change difference in the annotated proteins between OSKM-expressing cells and control cells.

In order to validate the transcriptomic analyses at the protein level we used proteome profiler human and mouse apoptosis arrays. The results show the overexpression of common apoptotic factors in both A549-rtTA-OSKM cells (e.g. TRAIL R1/R2, FAS or phospho-p53) and L1475luc-OSKM cells (e.g. cleaved-caspase 3 or cytochrome 3), as well as the downregulation of anti-apoptotic factors (e.g. XIAP), upon doxycycline exposure (**Fig. 3c, d** and **Supplementary Fig. 4d-g**). Moreover, increased levels of p-p53/p53, p27, and HSP60 in human and/or lung cancer cell lines are consistent with the implementation of senescence programmes (**Fig. 3c, d**).

Overall, transcriptomic and proteomic analyses are consistent with the induction of apoptosis and senescence upon OSKM expression in both human and mouse lung cancer cell lines.

### OSKM-induced apoptosis and senescence are partially driven by p21

One of the most interesting genes identified on our analyses was *CDKN1A*, coding for the CDK inhibitor p21. This protein has been described as being involved both in apoptosis and cell senescence^29^. We confirmed the upregulation of p21 at protein levels after induction of OSKM expression in both, human and mouse cells (**Fig. 4a**). To functionally address the involvement of p21 on the effect observed after expressing the reprogramming factors on cancer cells, we decided to reduce its expression by using an shRNA targeting *CDKN1A*. Lentiviral transduction of the A549-rtTA-OSKM cells with this plasmid caused a reduction on p21 mRNA and protein levels (**Fig. 4b**).

**Figure 4.**
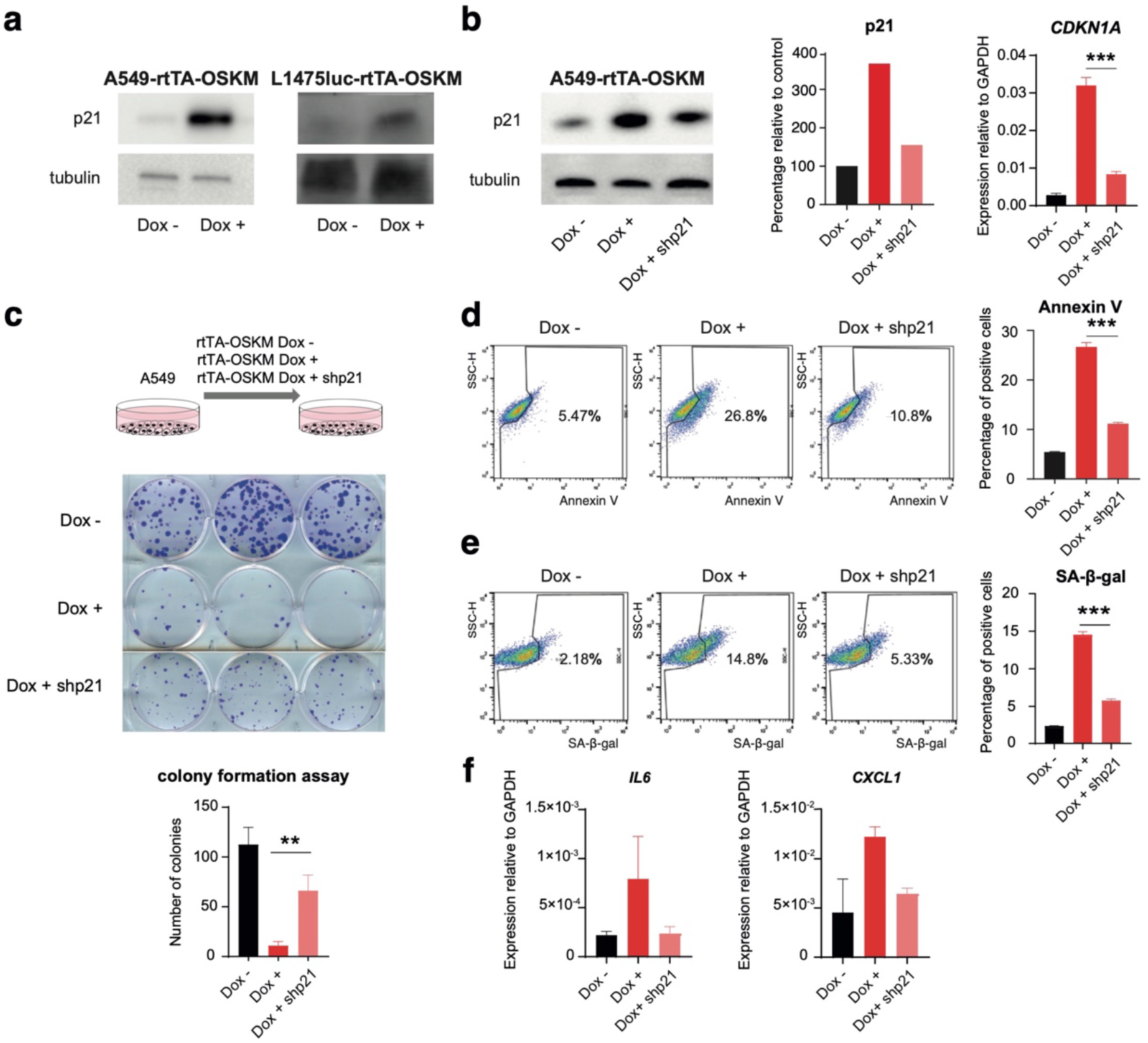
OSKM induction of cell apoptosis and senescence is mediated by p21. **(a)** Western blot of p21 in A549-rtTA-OSKM cells (left) and L1475luc-rtTA-OSKM cells (right) expressing (Dox +) or not (Dox -) the reprogramming factors. **(b)** Expression of p21 by Western blot (left panel, blot and quantification) and RT-qPCR (right panel) in A549-rtTA-OSKM cells expressing (Dox +) or not (Dox -) the reprogramming factors, and in cells expressing the factors and after knockdown of p21 (Dox + shp21). **(c)** Schematic representation of the experimental system used to test the proliferation of A549-rtTA-OSKM cells after knockdown of p21 (top panel). Representative images of a clonal expansion assay (middle panel) and crystal violet staining quantification (bottom panel) for the indicated experimental conditions. (**d**) Annexin V and (**e**) SA-ß-gal activity measurements by flow cytometry (left) and quantification (right) for the indicated experimental conditions. (**f**) Expression levels by RT-qPCR of *IL6* and *CXCL1* mRNA. Statistical significance was calculated using Student’s t-test, ***P<0.001; **P<0.01; *P<0.05. Data are mean ± SD.

We then overexpressed the reprogramming factors alone or in combination with the shRNA targeting p21 and assessed the growth of these cell cultures after 14 days. Cells with reduced p21 showed a partial recovery of cell growth (**Fig. 4c**), and this was not caused by an altered expression of OSKM since mRNA levels of the cassette were not reduced (**Supplementary Fig. 5a**). To further assess the impact of reducing p21 levels on the context of expression of OSKM, we analyzed apoptosis and senescence induction of cells lentivirally-transduced with the shRNA targeting *CDKN1A*. Consistent with **Fig 4c**, flow cytometry analysis of Annexin V expression showed a partial rescue of OSKM-induced apoptosis when p21 levels were reduced (**Fig. 4d**). Similarly, the shRNA against *CDKN1A* caused reduced induction of cell senescence by OSKM, as judged by SA-ß-gal staining (**Fig. 4e**), and a reduction of the expression of some genes encoding for common SASP markers (**Fig. 4f**).

Together, these data point to p21 as a relevant player mediating the induction of apoptosis and senescence by OSKM expression in cancer cells. Reducing p21 induction leads to a partial recovery of the growth of A549 lung cancer cells. However, this recovery was not sufficient to allow full cell reprogramming by OSKM, since we never observed the emergence of iPSC colonies from these cell cultures (**Supplementary Fig. 5b**). This was not due to residual expression of p21 since complete CRISPR-mediated deletion of *CDKN1A* did not result either in iPSC colony formation not even when combined with a pan-Caspase inhibitor (**Supplementary Fig. 5c**).

### Oct4 and Klf4 are the main factors contributing to apoptosis and senescence induction

Next, we explored the individual contribution of each reprogramming factor to the impaired growth caused by OSKM expression in lung cancer cells. To address this, we decided to individually express each factor and compare the growth of these cells with the ones expressing the polycistronic cassette (**Fig. 5a**). We confirmed that the expression of each individual factor reached similar levels to the ones obtained with the polycistronic OSKM cassette (**Supplementary Fig. 6a**). After 3 days in culture, we clearly observed that cells expressing OCT4 or KLF4 showed a reduced cell growth comparable to the one observed when we expressed the 4 factors (**Fig. 5b**). Longer cell culture times confirmed that the individual expression of OCT4 or KLF4 produced a lower number of cell foci similar to the one obtained with the combined expression of OSKM (**Fig. 5c**).

**Figure 5.**
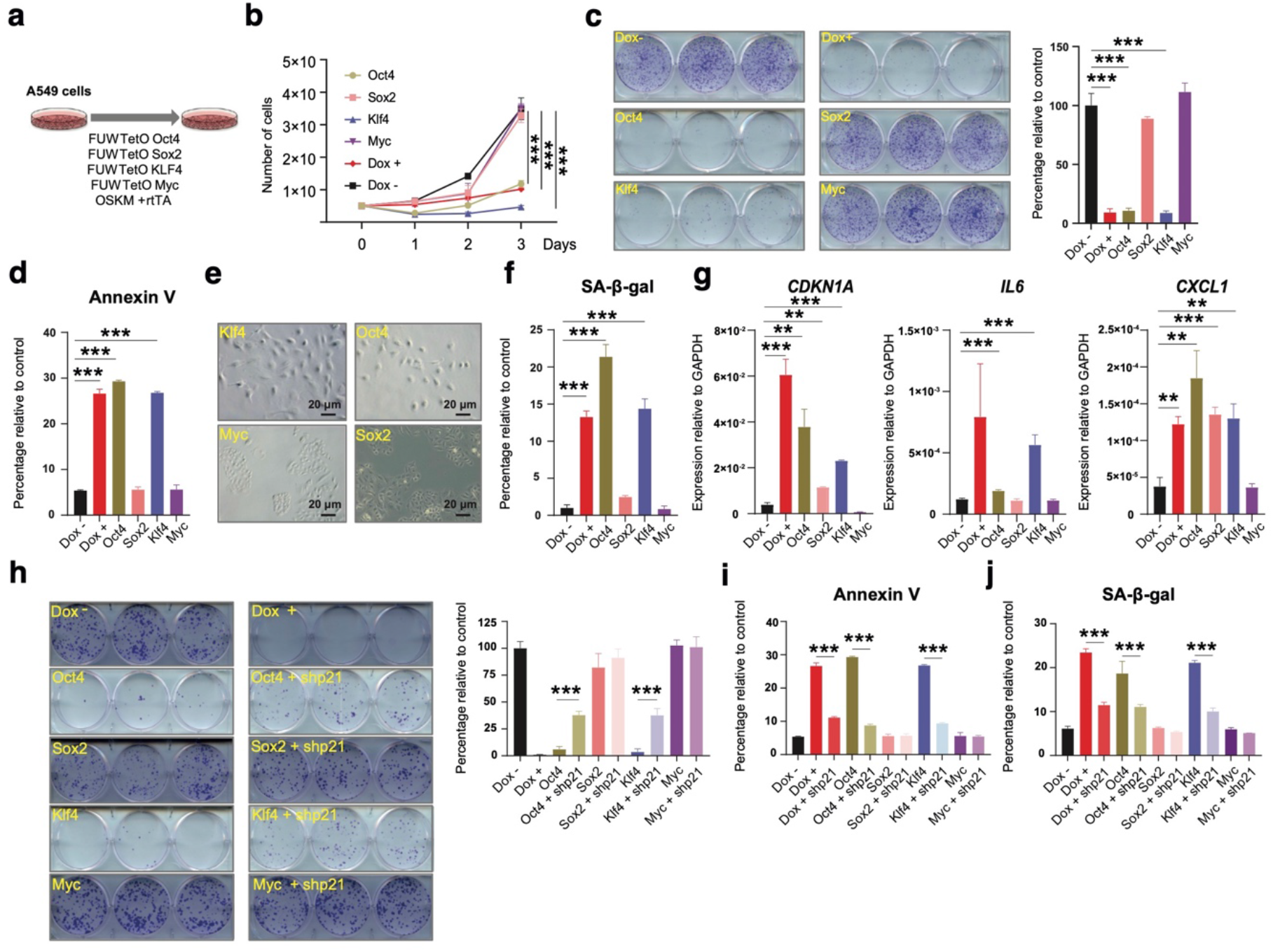
Contribution of individual reprogramming factors to the impaired cell growth. **(a)** Schematic representation of the experimental system used. **(b)** Analysis of the proliferative capacity and **(c)** Clonal expansion assay and quantification of A549 cells expressing individual reprogramming factors compared to A549-rtTA-OSKM cells expressing (Dox +) or not (Dox-) the four Yamanaka factors. **(d)** Quantification of Annexin V levels of cells expressing factors as in (b) measured by flow cytometry. **(e).** Representative light microscopy images of A549 cells expressing individual reprogramming factors. **(f)** Measurement of SA-ß-galactosidase activity of cells expressing factors as in (b) by flow cytometry. **(g)** Expression levels by RT-qPCR of *CDKN1A*, *IL6* and *CXCL1* in cells expressing factors as in (b). **(h)** Clonal expansion assay and quantification of A549 cells expressing individual reprogramming factors alone or after p21 knockdown (shp21) compared to A549-rtTA-OSKM cells expressing (Dox +) or not (Dox -) the four Yamanaka factors. **(i)** Quantification of Annexin V levels of cells expressing factors as in (h). **(j)** Measurement of SA-ß-galactosidase activity of cells expressing factors as in (h) by flow cytometry. Statistical significance was calculated using Student’s t-test, ***P<0.001; **P<0.01; *P<0.05. Data are mean ± SD.

Since we detected that apoptosis and senescence induction were contributing to the reduced growth of lung cancer cells triggered by OSKM expression, we evaluated how the individual factors were affecting these processes. Flow cytometry analysis of Annexin V showed that OCT4 and KLF4 induced apoptosis at a level similar to the combined expression of the 4 factors, OSKM (**Fig. 5d**). The microscopic observation of these cell cultures expressing the individual factors revealed that OCT4 and KLF4 expression caused a change in cell morphology that resembled the phenotype of senescent cells (**Fig. 5e**). In agreement, measurement of SA-ß-gal activity confirmed that OCT4 and KLF4 expression induced this marker of cell senescence (**Fig. 5f**), as well as an increase of mRNA levels of *CDKN1A* and other genes involved in this process (**Fig. 5g**).

To evaluate whether OCT4 and KLF4 were acting through the induction of p21, similarly to what we had already observed when we were using the 4 factors OSKM, we combined the expression of each individual factor with the expression of the shRNA targeting *CDKN1A*. First, we observed that individual OCT4 and KLF4 were both increasing p21 to similar levels to OSKM (**Supplementary Fig. 6b**). Also, we confirmed that our shRNA targeting *CDKN1A* was capable of reducing the expression of p21 (**Supplementary Fig. 6b**). Analysis of cell growth at long periods of time showed that reducing the expression of p21 in the context of single overexpression of OCT4 or KLF4 resulted in the partial protection of the growth of lung cancer cells (**Fig. 5h**). This alleviation of the reduced growth induced by OCT4 and KLF4 after reducing the expression of p21 was also accompanied by a partial reduction in apoptosis (**Fig. 5i**; **Supplementary Fig. 6c**) and cell senescence (**Fig. 5j; Supplementary Fig. 6d**).

In summary, the expression of OSKM did not promote efficient cellular reprogramming of lung cancer cells, but impaired growth by engaging apoptosis and senescence. OCT4 and KLF4 seemed to be the most relevant factors responsible of this effect and p21 was an important mediator in this process.

### OSKM expression limits tumor initiation and progression in mouse models of transplanted lung cancer cells

The aforementioned *in vitro* experiments using human A549 and mouse L1475luc lung cancer cells showed that OSKM expression impairs cell proliferation and growth by the induction of apoptosis and cellular senescence. To validate *in vivo* the impact of OSKM expression on lung tumor cell growth, we first subcutaneously transplanted L1475luc-rtTA cells (left flanks, TA) and L1475luc-rtTA-OSKM cells (right flanks, 4F) in C57BL/6 mice. In our experimental settings doxycycline (0.2 or 1 mg/mL) was uninterruptedly provided in the drinking water from lung cancer cell transplantation to the experimental end point, and the tumor burden was longitudinally monitored by measuring luciferase activity with an IVIS bioluminescence imaging system (**Fig. 6a**). As shown in **Fig. 6b**, at 10 days post-transplantation, a significant proportion of cells expressing OCT4, SOX2 or MYC were detected in L1475luc-rtTA-OSKM tumor sections (Oct4^+^ 7.701±2.727%; Sox2^+^ 3.108±2.191%; c-Myc^+^ 24.96±2.662%) relative to control L1475luc-rtTA tumor sections (Oct4^+^ 0.648±0.149%, p = 0.011; Sox2^+^ 0.189±0.193%, p = 0.08; c-Myc^+^ 14.15±4.972%, p = 0.029, relative to OSKM-expressing cells), where the levels of reprogramming factors remained essentially undetectable (**Fig. 6c**). Early post-transplantation times (from day 1 to day 12) presented a similar tumor progression for both L1475luc-rtTA cells and L1475luc-rtTA-OSKM cells (**Fig. 6d, e**; **Supplementary Fig. 7a**). However, OSKM-expressing tumors significantly reduced their growth rate from day 12 to day 26 post-transplantation when compared to L1475luc-rtTA tumors, thereby resulting in a reduced tumor burden (**Fig. 6d, e**; **Supplementary Fig. 7a**). Similar results were obtained using two different doses of doxycycline to induce OSKM expression (**Fig. 6e**; **Supplementary Fig. 7a**). Of note, this reduction in the luciferase signal inversely correlates with a significant increase in apoptosis markers in L1475luc-rtTA-OSKM cells (cleaved caspase 3^+^ ratio to tumor area 0.215 ±0.109) relative to L1475luc-rtTA cells (cleaved caspase 3^+^ ratio to tumor area 0.078±0.033%, p < 0.0001) (**Fig. 6f, g**) concomitant with a trend of higher p21 expression in L1475luc-rtTA-OSKM cells (p21^+^ 0.694±0.227%) relative to L1475luc-rtTA cells (p21^+^ 0.579±0.184%, p = 0.138) (**Fig. 6h, i**). Tumor areas enriched in cleaved caspase 3^+^ cells and cells expressing p21correlate with the accumulation of cells expressing Oct4, Sox2 and c-Myc (**Supplementary Fig. 7b**).

**Figure 6.**
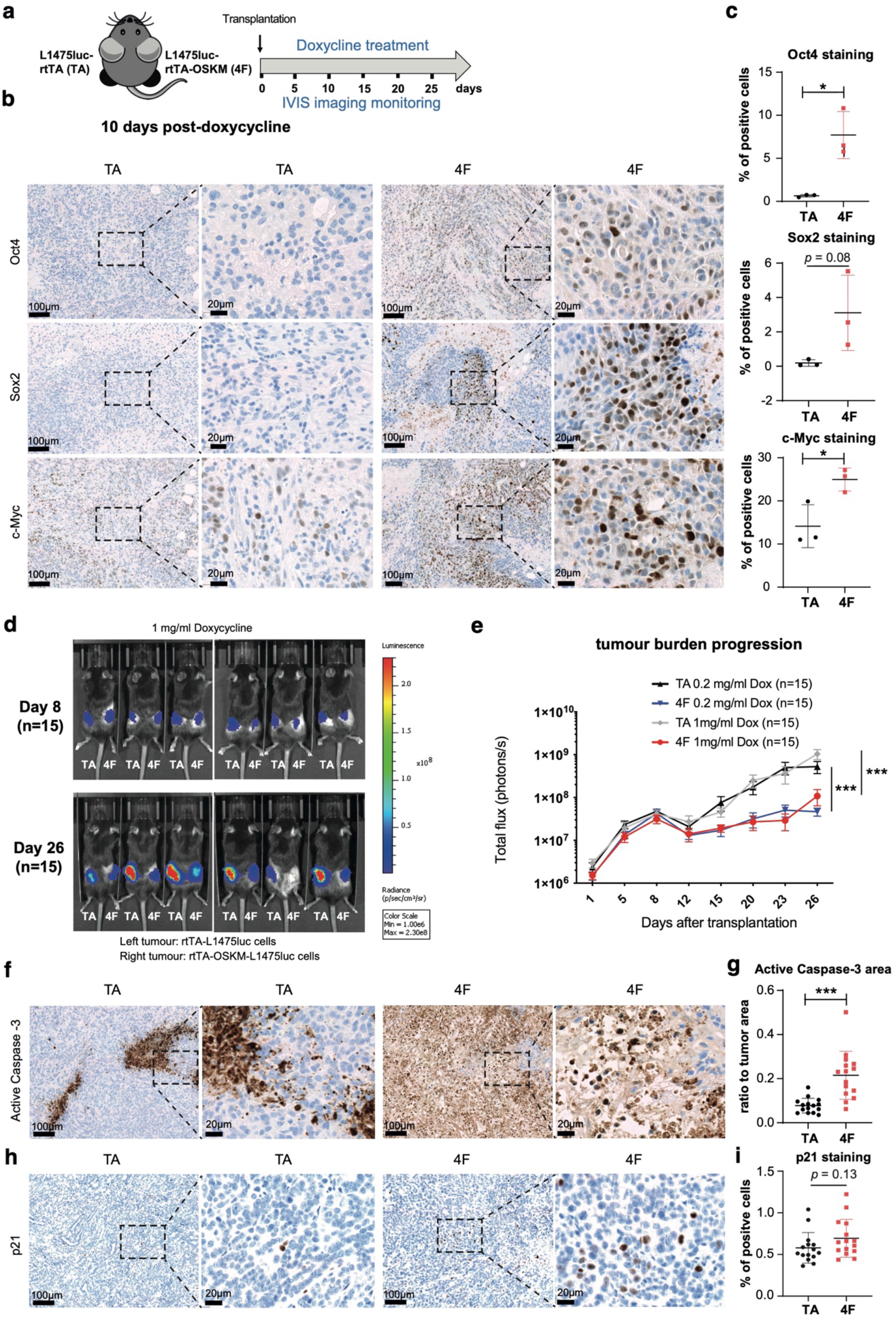
Expression of OSKM impairs growth of subcutaneous tumors. **(a)** Schematic representation of the in vivo experimental system. L1475luc-rtTA cells were injected in the left flanks (TA) and L1475luc-rtTA-OSKM cells in the right flanks (4F) of C57BL/6 mice that were treated with doxycycline and monitored for the indicated period of time (n=15). **(b)** Immunohistochemistry and **(c)** quantification of the expression of Oct4, Sox2 and c-Myc at 10 days post-doxycycline administration in tissue sections from TA and 4F tumors. Dashed lines indicate magnified areas. **(d)** Representative images of TA and 4F tumors exhibiting luciferase activity measured by IVIS at the indicated time points after transplantation. **(e)** Longitudinal quantification of TA and 4F tumor growth in mice treated with 0.2 mg/ml and 1 mg/ml doxycycline. **(f)** Immunohistochemistry and **(g)** quantification of the expression of active Caspase-3 at the end point (26 days) in tissue sections from TA and 4F tumors. Dashed lines indicate magnified areas. **(h)** Immunohistochemistry and **(i)** quantification of the expression of active p21 as in (f) and (g). Statistical significance of the different tumor growth was analyzed by two-way ANOVA, ***, p<0.001. For the histological analysis, the Student’s t-test was used, ***P<0.001; **P<0.01; *P<0.05. Data are mean ± SD.

We next sought to ascertain the effect of OSKM expression in lung cancer L1475luc cells in an orthotopic model where the cells were transplanted in the lungs of C57BL/6 mice via tail-vein injection. Similarly to the allograft experiments, 1 mg/mL doxycycline was provided in the mouse drinking water from initial cell transplantation times to the end point (for a total of 2 weeks), and the tumor burden was longitudinally monitored by using an IVIS bioluminescence imaging system (**Fig. 7a**). While the luciferase activity of both L1475luc-rtTA cells (TA) and L1475luc-rtTA-OSKM cells (4F) was shown to be comparable at day 7 post-transplantation, the tumor progression was drastically diminished at day 14 post-transplantation in OSKM-expressing cells (119,154±56,871 photon/s in L1475luc-rtTA-OSKM cells relative to 667,055±285,305 photon/s in L1475luc-rtTA cells, p = 0.0191) (**Fig. 7b, c**; **Supplementary Fig. 8**). Accordingly, lung histology sections from L1475luc-rtTA-OSKM transplanted mice showed positive intratumoral levels of cleaved caspase 3 activation, and evidence of p21 and Oct4 expression (**Fig. 7d,e**).

**Figure 7.**
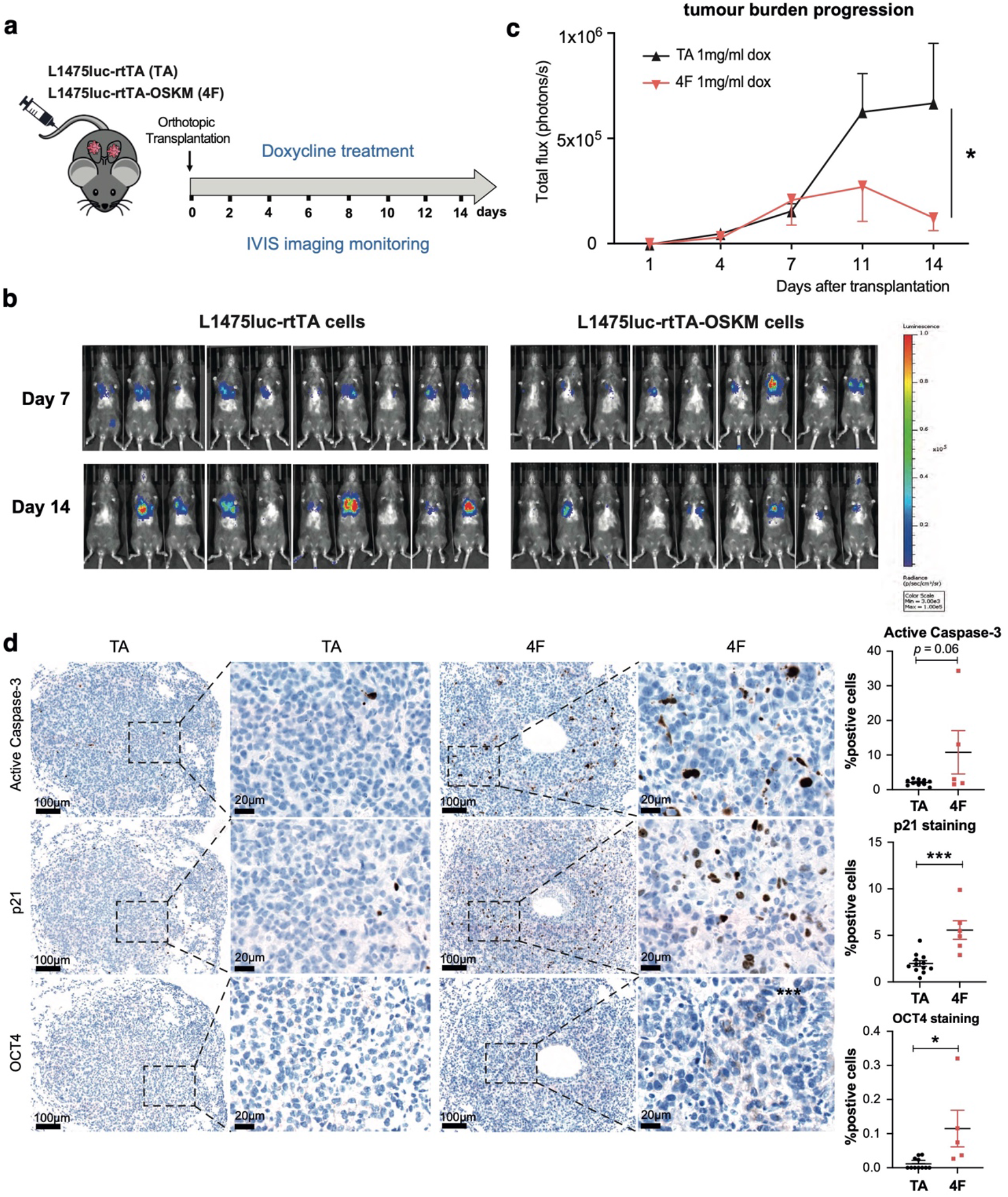
Expression of OSKM impairs lung tumor growth of orthotopically transplanted cells. **(a)** Schematic representation of the in vivo experimental system. L1475luc-rtTA (TA) or L1475luc-rtTA-OSKM (4F) cells were tail-vein injected for orthotopic transplantation in the lungs of C57BL/6 mice that were treated with doxycycline and monitored for the indicated period of time (n=9-10). **(b)** Representative images of TA and 4F tumors exhibiting luciferase activity measured by IVIS at the indicated time points after transplantation. **(c)** Longitudinal quantification of TA and 4F tumor growth in mice treated with 1 mg/ml doxycycline. **(d)** Representative images (left) of immunohistochemistry for active Caspase-3, p21 and Oct4 in TA mice and 4F mice and quantifications (right). Statistical significance of the different tumor growth was analyzed by two-way ANOVA, *, p<0.05. For the histological analysis, the Student’s t-test was used, ***P<0.001; **P<0.01; *P<0.05. Data are mean ± SD.

Taken together, tumor allografts and orthotopic transplantation models of murine lung cancer L1475luc cell lines confirmed the induction of apoptosis upon OSKM expression correlating with a reduction of the tumor burden. These results indicate that OSKM expression in lung cancer cells limits tumor initiation and progression *in vivo*, thereby validating our *in vitro* findings.

### Expression of OSKM factors results in reduced tumor burden in a GEMM of KRAS-driven lung cancer

Finally, we aimed to validate our observations using a distinct and more physiologically relevant murine model of KRAS-driven lung adenocarcinoma induced by endogenous expression of oncogenic KrasG12V. For this purpose, we made use of the *Kras-FSFG12V* model, in which the expression of oncogenic KrasG12V is prevented by a *frt*-flanked DNA sequence, which allows for lung tumor induction following intra-tracheal administration of adenovirus-FLP (AdFLP)^30^. *Kras-FSFG12V* mice were crossed with reprogrammable i4F-B mice^31^, carrying the transcriptional activator (rtTA) within the ubiquitously-expressed *Rosa26* locus and a single copy of a lentiviral doxycycline-inducible polycistronic cassette encoding the four murine Yamanaka factors *Oct4*, *Sox2*, *Klf4* and *c-Myc*^31^, leading to *Kras^FSFG12V^*^/*+*^;*i4F^KI/WT^* mice, from here onwards KrasG12V;OSKM mice (**Fig. 8a**). To assess the effect of OSKM expression in murine lung tumorigenesis, we treated KrasG12V and KrasG12V; OSKM mice with doxycycline in the drinking water at 1 mg/mL, for 1 week/month during 3 consecutive months, and starting at 6 months after intra-tracheal administration of AdFLP (**Fig. 8a**). KrasG12V mice present at this stage (6 months) varying degrees of tumor development, allowing for the observation of abundant early lung hyperplasias and adenomas, with occasional progression to adenocarcinomas^32^. As an internal control of OSKM induction, KrasG12V; OSKM mice and their control counterparts were treated for one week with 1 mg/mL doxycycline in drinking water, culled, and lung homogenates were subjected to RT-qPCR analyses that confirmed the upregulation of *Oct4*, *Sox2* and *Klf4* genes (**Supplementary Fig. 9a**).

**Figure 8.**
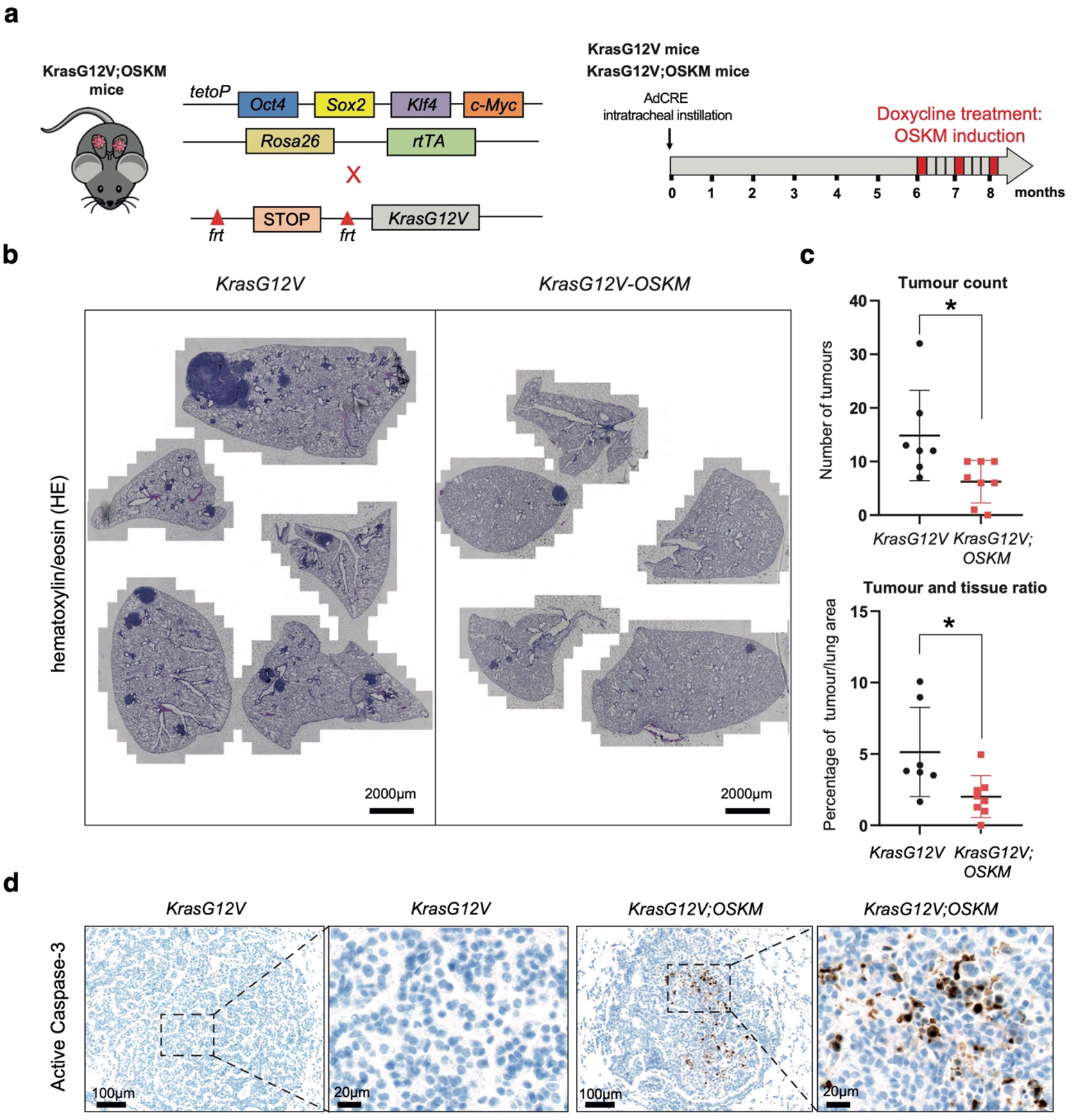
OSKM expression reduces lung tumor burden in a KrasG12V mouse model. **(a)** Schematic representation of the generation of the genetically-engineered mouse model. Inducible OSKM expressing mice were crossed with KrasG12V mice to generate KrasG12V;OSKM mice (left). Experimental design summary. Experimental mice (KrasG12V n=7 and KrasG12V;OSKM n=8) were intranasally-treated with adenovirus-FLP (AdFLP) expressing flippase recombinase to activate oncogenic KrasG12V mutation. Mice were treated with 1mg/ml doxycycline (one week on, three weeks off, during 3 consecutive cycles) in drinking water from 6-month post lung cancer initiation. Mice were culled after the last one-week long doxycycline treatment and the lungs were collected for histology analyses (right). **(b)** Representative H&E staining images of lung sections from KrasG12V (left) and KrasG12V;OSKM mice (right), as indicated. **(c)** Number of tumors per mouse (top graph) and percentage of tumor per lung are (bottom graph) in the indicated experimental groups. Each data point represents one mouse. **(d** Representative images of immunohistochemistry for active Caspase-3 in lungs of KrasG12V mice (left) and KrasG12V;OSKM mice (right), as indicated. Statistical significance was calculated using Student’s t-test, ***P<0.001; **P<0.01; *P<0.05. Data are mean ± SEM.

Given the limited number of mice included in the experiment (n=8 for KrasG12V;OSKM and n=7 for KrasG12V), we only collected lung samples at the endpoint (after the third weekly doxycyline treatment at 8 months post-intratracheal injection). This precluded us from performing a thorough histological analyses at earlier timepoints. However, some tumors from KrasG12V; OSKM mice showed cleaved caspase 3 positivity while this was never observed in KrasG12V mice (**Fig. 8d**). Regarding p21, no differences were observed between the two experimental groups at the endpoint (**Supplementary Fig. 9b**). Remarkably however, three consecutive cycles of OSKM induction by doxycycline administration resulted in a significant reduction of the tumor burden as per the number of tumors (6.250±3.955 tumors in KrasG12V;OSKM mice relative to 14.86±8.345 tumors in KrasG12V mice; p = 0.225) and percentage of tumors/lung area (2.011±1.465% in KrasG12V;OSKM mice relative to 5.135±3.121% in KrasG12V mice) (**Fig. 8b and c**).

Collectively, data obtained using allografts, orthotopic transplantation and genetically-engineered KRAS-driven lung cancer mouse models clearly demonstrate that transient OSKM expression impairs tumor initiation and progression and results in a significant reduction of the tumor burden.

## Discussion

Transcription factor-mediated reprogramming to pluripotency is an ideal in vitro system to study cellular processes involving changes in cell identity^3^. Pluripotency and cancer share many commonalities, such as similar transcriptional signatures, and are the result of two analogous processes, reprogramming and transformation, respectively^12^. Despite these similarities, and the fact that reprogramming factors have been individually involved in cancer, there is a striking lack of reports showing a full and efficient reprogramming of cancer cells to pluripotency. Apart from a few descriptions of successful reprogramming using blood cancers and some metastable states achieved in epithelial cancer cells^33^, it is widely accepted that epithelial tumors, and specifically lung cancer cells, are highly resistant to reprogramming to pluripotency. To gain insights into the molecular mechanisms underlying their resistance to reprogramming, we have expressed the Yamanaka factors in human and mouse lung cancer cells, and examined the effect on in vitro cell growth and in a variety of murine lung cancer models.

First, we observed that cultures expressing OSKM factors were impaired in cell growth due to a combination of apoptosis and senescence induction. This was shown by cell proliferation and functional analyses, and further confirmed using high throughput transcriptomic and proteomic approaches. Given the potential role of the reprogramming factors as oncogenes, it is tempting to speculate that this response resembles oncogene-induced apoptosis and senescence^34^. In fact, apoptosis and senescence induced by the expression of OSKM is known to limit the efficiency of reprogramming to pluripotency^13–18^. A similar situation has been previously described for leukemia, in which OSKM expression results in an epigenetic remodeling leading to increased chromatin accessibility near genes encoding pro-apoptotic regulators^35^.

When we expressed the individual Yamanaka factors, we observed that single expression of Oct4 or Klf4 was sufficient to recapitulate the effects obtained with the full OSKM cassette. In line with this, Oct4 has been shown to activate Caspase 3 and 8 during cell reprogramming^36^, while Klf4 has been reported to regulate adult lung tumor-initiating cells and represses K-Ras-mediated lung cancer^37^. Interestingly, Sox2 and Klf4 have been pointed out as the relevant reprogramming factors inducing leukemic apoptosis, exemplifying the context dependent effect of the Yamanaka factors in cancer cells of different origin.

In our work, besides apoptosis, a distinctive feature of OSKM expression in lung cancer cells is the induction of cell senescence. A key gene involved in both senescence and apoptosis, is the cell cycle and CDK inhibitor p21^29^. Remarkably, we observed an increased expression of p21 both at the mRNA and protein levels in lung cancer cells after activation of the OSKM cassette or after individual expression of Oct4 or Klf4. This expression correlates with the induction of apoptosis and senescence, and it is absent when the other two factors, Sox2 and c-Myc, were expressed. In agreement, knockdown of p21 expression using shRNA, alleviates the apoptosis and senescence induction and partially rescues the growth of full OSKM or individual Oct4 or Klf4 expressing lung cancer cells. In this regard, mechanistically, Klf4 has been previously shown to regulate the levels of p21 in a cancer context dependent manner^9^. Together, these results causally connect OSKM-mediated induction of p21 with the implementation of apoptosis and senescence during failed reprogramming of lung cancer cells. However, CRISPR-mediated knockout of *CDKN1A* (encoding p21) in lung cancer cells is not sufficient to render them susceptible to OSKM-mediated reprogramming to iPSCs, even when combined with a pan-caspase inhibitor that also partially rescues the observed phenotypes.

To validate these results in vivo, we used three murine models of carcinogenesis, namely subcutaneous tumor allograft implantation, orthotopic lung transplantation, and genetically engineered *Kras*-driven lung cancer development. Consistently, our experiments show that OSKM expression in lung cancer cells result in impaired tumor growth concomitant with an accumulation of p21 and the induction of apoptosis.

Our previous work had already shown that neoplastic transformation and pluripotency are two incompatible cellular fates^25^. Cells expressing OSKM trigger a molecular trajectory that, although initially intersecting with that originated by the transforming activity of Ras/c-Myc, diverges towards an independent cellular state^26^. Identifying the key regulatory elements of cancer cell identity is instrumental for the discovery of novel vulnerabilities that would allow us to design new and more effective therapeutic strategies^4,38,39^.

## Methods

### Cell line generation

L1475luc cells are a kind of gift from Dr. Carla P Martins (AstraZeneca, Cambridge, UK)^27^. These cells were cultured in DMEM/F12 medium with 10% FBS. Transposon vectors expressing OSKM and rtTA are kind gifts from Professor Keisuke Kaji, PiggyBac vector was purchased from System Biosciences (PB210PA-1; Palo Alto, CA). For transfection, 0.5×10^6^ cells were seeded in 1 well of a 6-well plate the night before. For each well, the following plasmid mixture solutions were added: 0.5 µg TetO-OSKM-mOrange, 0.5 µg rtTA-puro and 1µg of PiggyBac (for OSKM expressing cells) or 0.5 µg rtTA-puro and 1 µg of PiggyBac (for control rtTA cells) in 100 µl Opti-MEM solution (31985062, Life Technologies) containing 5 µl of X-tremeGENE 9 transfection reagent (XTG9-RO Roche, Sigma-Aldrich). Three days after transfection, puromycin (A1113803, Life Technologies) was added to transfected cells. After 7 days, to select for the transfected cells, doxycycline (D9891, Sigma-Aldrich; 1 µg/ml final concentration) was added for overnight treatment to induce mOrange reporter gene expression. The next day, mOrange positive cells were sorted in BD Influx cell sorter (BD Biosciences). Sorted cells were maintained in DMEM/F12 medium with 10% FBS containing puromycin. All cells were used before passage 10 in all the experiments. In the experiments with A549 cells, we introduced the FUW-M2rtTA construct^40^ alone or in combination with Tet-O-FUW-OSKM^41^, TetO-FUW-OCT4 (Addgene #20323) TetO-FUW-Sox2 (Addgene #20326) TetO-FUW-Myc (Addgene #20324) or TetO-FUW-Klf4 (Addgene #2032)^42^. To induce the expression of the reprogramming factors, cells were treated with doxycycline at 1 µg/mL (Sigma-Aldrich) and the culture medium was renewed every 2 days during the treatment. All the cell lines used were maintained in fibroblast medium: DMEM (Dulbecco’s Modified Eagle Medium) with high concentration of glucose (4500 mg/L) (Sigma-Aldrich), 10% FBS (Sigma-Aldrich), 1% penicillin/streptomycin (Sigma-Aldrich), 1% glutamine (Sigma-Aldrich). All lines were kept in an incubator at 37°C and 5% CO2 atmosphere. To induce OSKM expression in L1475luc-OSKM cells, doxycycline was added (D9891, Sigma-Aldrich; 1 µg/ml final concentration except otherwise stated). All cells used were routinely tested for mycoplasma.

### Cell proliferation and apoptosis

For cell proliferation, A549-rtTA-OSKM or L1475luc-OSKM cells were seeded in 6-well plates (0.5×10^6^ cells/well). Doxycycline was added to each group (n=3 wells/group) on the treatment plate and a control plate without doxycycline was also prepared. We analyzed the proliferative capacity during the first 3 days by counting cells on a Neubauer chamber (Hausser Scientific). For real-time live imaging, plates were placed in Incucyte Zoom System (Essen BioScience) and images were taken every 3 h for about 120 h, at 16 field/image, with a 10x objective. To detect apoptotic cells, Incucyte Caspase-3/7 Green Dye (4440, Essen BioScience) was added, and images were taken in both, bright field (for cell proliferation) and green fluorescence channel for Alexa 488 (for apoptosis detection). Data analysis was performed with the Incucyte Zoom software (2016B). Cell confluence was calculated by label free identification of cell and blank areas. Cell apoptosis was calculated by measuring green pixels per area. The experiments were performed in 3 biological replicates.

Cell proliferation and apoptosis were also determined by flow cytometry. For L1475luc-OSKM, cells were studied daily from day 1 to day 5. Cell number was calculated with reference beads (424902, BioLegend, using 50 µl beads per 500 µl cell suspension) and apoptosis was detected with eBioscience Annexin V Apoptosis Detection Kit (88-8005-74, ThermoFisher). For apoptosis, DAPI was used to label dead cells. Cell acquisition was done in BD LSRFortessa Cell Analyzer. Flow cytometry data were analyzed with FlowJo V10.7 software, and apoptotic cells were defined as DAPI^-^ Annexin V^+^ cells. In A549, for the detection of cleaved Caspase 3 we used the PE active Caspase 3 apoptosis kit (550914, BD Biosciences) and for the detection of Annexin V we used the FITC Annexin V Kit (Immunostep). We analyzed 1×10^4^ cells in each sample with a FACScan cytometer (BD Biosciences).

To inhibit the apoptosis process, we used the inhibitor Z-VAD-(Ome)-FMK (MedChem) at a concentration of 20 µM. The inhibitor was renewed every 2 days during the treatment.

### Clonogenicity assay

For the clonogenicity assays, we seeded 500-1000 cells expressing or not the reprogramming factors in 6-well plates, in triplicate, for each condition. The medium and doxycycline, in the corresponding condition, were renewed every 2 days. After 10-14 days we evaluated cell growth and foci formation with crystal violet staining. Where indicated, crystal violet was eluted with 10% acetic acid, and the optical density was measured by spectrophotometry (at 570 nm) or colonies were counted either with ImageJ or software8.

### Colony formation assay in soft agar

6-well plates were coated with a solid base of 1 mL of agar (0.8% in complete DMEM) (Thermo Fisher). On top of this solid base, 2.5×10^3^ A549 cells were seeded on 1 mL of agar (0.35% in fibroblast complete DMEM). Fresh medium and doxycycline were added at a concentration of 1 µg/ml, every 2 days for 10 days, and the number of colonies per well was quantified using an AxioVert.A1 Microscope (Zeiss). To detect apoptosis in soft agar colonies, we added cellEvent™ Caspase-3 Green detection reagent (C10423, Invitrogen) at 2 µM.

### Apoptosis protein array

For the apoptosis protein array assay, the kits used were: Proteome Profiler Human (ARY009) and Mouse (ARY131) Apoptosis Array Kits, following the manufacturer’s specifications.

## RNA-seq

RNeasy Mini Kit (Qiagen 74104) was used for total RNA extraction on sub-confluent cells in four replicates according to the manufacturer’s protocol. The quality and concentration were assessed with the Agilent RNA Nano 6000 kit (Agilent 5067-1511) on Agilent Bioanalyzer 2100 instrument. RNA-seq libraries were prepared by BGI. Sequencing reads were mapped to hg38 using RSEM and bowtie2. Differentially expressed genes were identified using DeSeq2^43^. Gene set enrichment analysis was performed using R packages ClusterProfiler^44^, Molecular Signature Database (MSigDB) Hallmarks gene set (Version 7.1.1), magrittr (Version 2.02).

### SA-**ß**-Gal activity

For the detection of SA-ß-Gal activity by flow cytometry we used the cellEvent™ Senescence Green Flow Cytometry Assay kit (C10840, Invitrogen). For the chemiluminescent detection of SA-ß-Gal activity, the Galacto-Light Plus™ beta-Galactosidase Reporter Gene Assay System kit (Applied Biosystems) was used, following the manufacturer’s instructions, except for the citric acid/sodium phosphate buffer that was used at pH 6.0, since it was better suited for the detection of this enzyme in the A549 cells analyzed.

### RT-qPCR

To measure gene expression, we extracted total RNA from cell cultures using the NucleoSpin® RNA Kit (Macherey-Nagel), following the manufacturer’s instructions, and converted it to cDNA using High-Capacity cDNA Reverse Transcription (Applied Biosystems). Quantification was performed using the NZYSpeedy qPCR Green Master Mix reagent (2X), ROX (NZYTech) and the AriaMx Real-Time PCR systems thermocycler (Agilent Technologies). For each reaction, 33 ng of cDNA, oligonucleotides at a final concentration of 0.25 µM, 5 µL of SYBR and nuclease-free water were used up to a final volume of 10 µl per reaction. GAPDH was used as housekeeping gene and the values of the analyzed genes were relativized to its expression level. Triplicates were used for each analysis. The results were analyzed with the AriaMx 1.0 software (Agilent Technologies) and the oligonucleotides used were purchased from Eurofins Genomics:

In the case of human oligonucleotides:

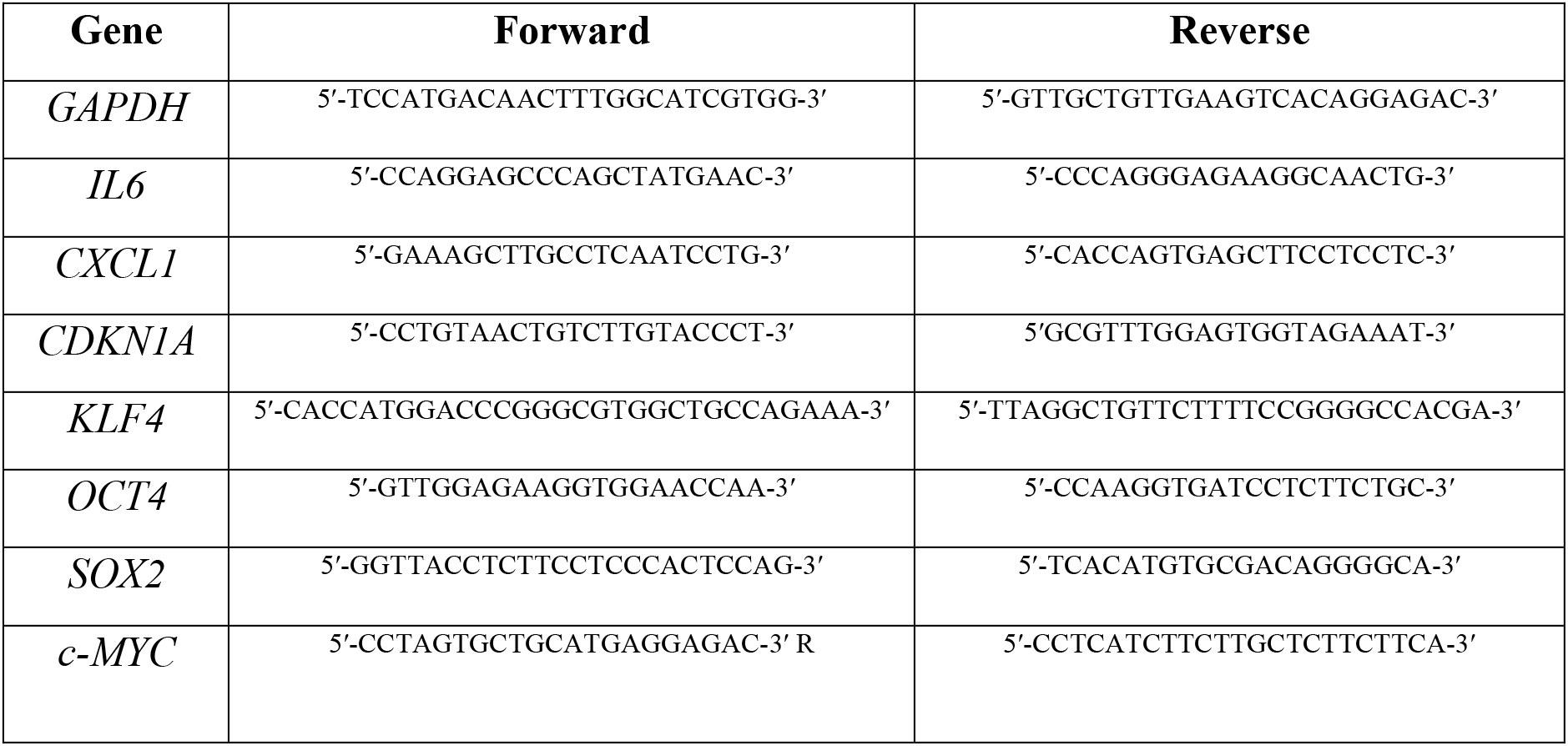

In the case of murine oligonucleotides:

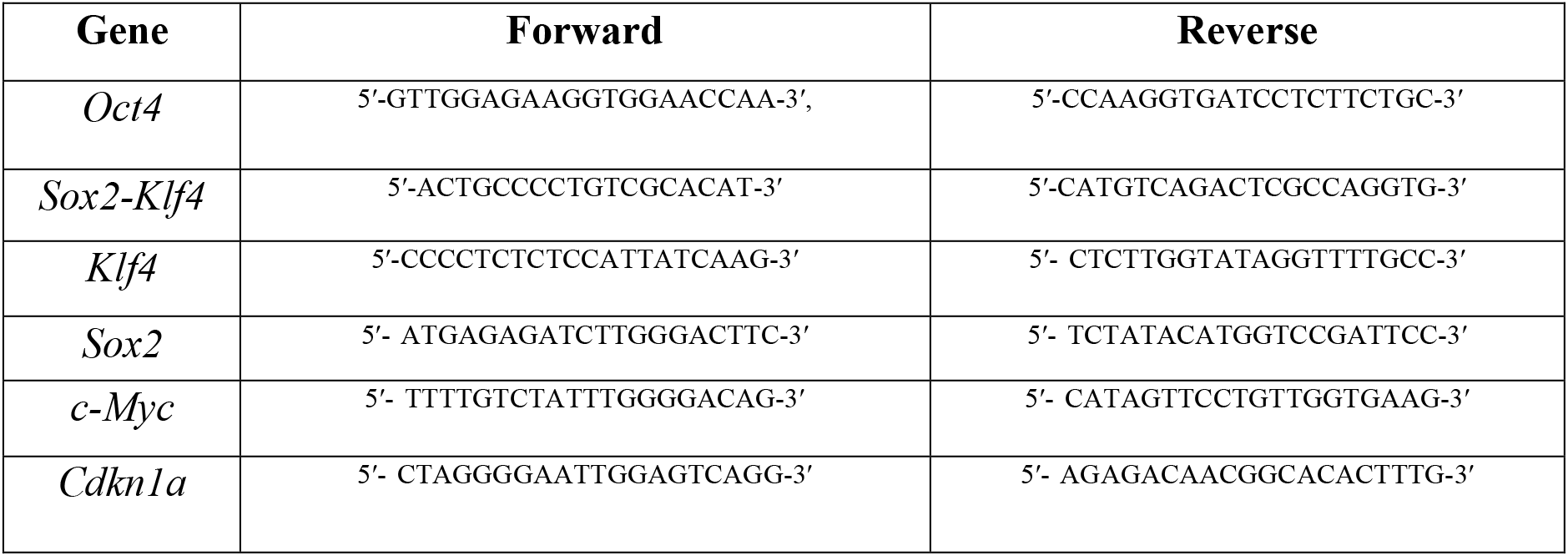

### Western blotting

Protein samples were prepared from cells lysed with RIPA buffer (R0278-50ML, Sigma-Aldrich) containing proteinase and phosphatase inhibitors (4693159001 and 4906845001, Sigma-Aldrich). Protein concentration was quantified with BCA assay (23225, Life Technologies). Precast gradient gels (4561086, Bio-Rad) were used to run the samples with 20-30 µg protein/well. After transfer and blocking with 5% milk solution, first antibodies (GAPDH: 10494-1-AP, Proteintech, 1:3,000; beta-actin: 4967, Cell Signaling Technology, 1:3,000; Oct4: 2840, Cell Signaling Technology, 1:100; Xiap: AF8221, Bio-Techne, 1:100; Beta-Tubulin: 2146, Cell Signaling Technology, 1:5,000; p21 Waf1/Cip1(12D1): 2947, Cell Signaling Technology, 1:1,000) were incubated overnight. Secondary antibodies from Li-Cor (Li-Cor Biosciences) were incubated for 1 h with 1:10,000 dilution. The membranes were imaged in a Li-Cor Odyssey device (Li-Cor Biosciences) or a ChemiDoc (Bio-Rad).

### Subcutaneous and lung orthotopic murine cancer models

Expression of mOrange was routinely examined in each batch of 4F cells and was always shown to be over 80% positive after doxycycline induction *in vitro*. Early passage (<10) 4F and TA cells were cultured until about 80% confluency before being detached by TrypLE Express enzyme (12604013, ThermoFisher). Cells were washed and resuspended in PBS and counted by Countess II automated cell counter (AMQA×1000, ThermoFisher) and over 90% cells were found to be viable. Female 8 weeks old C57BL/6 mice were purchased (Charles River Laboratories) and acclimatized to the local facility conditions at least 7 days before use. Mice were treated with doxycycline formulated in water (1 mg/ml with 7% sucrose or 0.2% mg/ml in 1.4% sucrose, as indicated) on day 0, irradiated with 4 Grays on day 1, and rested for 24 h before cell injection on day 2. TA or 4F cells (2×10^6^) were injected subcutaneously in 200 µl of DMEM at each flank of the mouse (left and right side, respectively). For orthotopic model, 2×10^5^ cells in 200 µl were intravenously injected into each mouse. Tumor bioluminescence signals were monitored twice per week using an IVIS Spectrum In Vivo Imaging System (PerkinElmer), at 10 min after injection of 200 µl/mouse, of 15 mg/ml D-luciferin (122799, PerkinElmer) solution in PBS. Exposure time was set to 15 s, 30 s, 1 and 3 min or optimal setting from the Living Image Software (V4.5.2, PerkinElmer). Non-saturated images (usually 30 s exposure for the orthotopic model) were quantified with the same software. In the orthotopic experiment, 2 mice were injected with PBS instead of cells and their bioluminescence signal was set as background value. The experiments were stopped after 26 days (in the subcutaneous model) or 14 days (in the orthotopic transplantation model) after cell injection. Subcutaneous tumor samples and lungs were collected in 10% formalin solution for histological analysis.

### KrasG12V-driven lung cancer model

KrasG12V mice, a kind gift from Professor Mariano Barbacid (CNIO, Madrid, Spain)^30, 32^ were crossed with OSKM mice, a kind gift from Professor Manuel Serrano (CNIO, Madrid, Spain)^31^, resulting in KrasG12V;OSKM mice. Transgenes were detected by PCR according to their published protocols. Mice were housed in groups under controlled conditions: 12 h light/darkness cycle at 21°C with free access to standard rodent chow and water. Both male and female mice (around 8-week-old) were used in the experiments. To induce KrasG12V expression, 2.5×10^7^ pfu adenovirus expressing flippase recombinase (Ad5CMVFlpo, VVC-U of Iowa-530, Viral Vector Core Facility, University of Iowa) was intranasally instilled into mice according to published protocol^30^. At 6 months post infection, we administered doxycycline in the drinking water for 3 cycles comprised of: 1 week doxycycline and 3 weeks resting with normal drinking water. After the last doxycycline treatment, animals were culled. Lung tissues were fixed in 10% formalin overnight, followed by ethanol treatment before paraffin embedding.

All experiments were approved for Ethical Conduct by the Home Office England and Central Biomedical Services (CBS), regulated under the Animals (Scientific Procedures) Act 1986 (ASPA), as stated in The International Guiding Principles for Biomedical Research involving Animals.

### Tissue histology and immunostaining

Formalin fixed, paraffin embedded blocks were cut into 5 µm tissue sections and used for histological analysis and immunostaining. Oct4, Sox2, p21, c-Myc and active Caspase-3 immunohistochemistry staining were performed by CNIO’s Histopathology Unit, and whole sections were scanned using an Axio Z1 slide scanner (Carl Zeiss). For quantification, tissue scans were analyzed with HALO software (V3.0.311.373, Indica Labs).

For tumor burden assessment in *KrasG12V;OSKM* mice, lung tissues were collected and fixed in 10% formalin before paraffin embedding. Paraffin blocks were serially cut into 5 µm sections (2-3 sections/slide), with 50 µm trimmed off every 10 slides, and at least 90 slides collected for each mouse. For H&E staining, 8-12 sections in regular intervals were used for each mouse and whole slides were scanned with Axio Z1 slide scanner (Carl Zeiss Ltd.). Tumor grading and counting were carried out according to a published protocol^45^. Briefly, tissues can be auto detected, and the regions of interest (ROI) corresponding to tumour areas were manually defined and analyzed by using HALO software (V3.0.311.373, Indica Labs). The ratio of tumor to lung area was calculated by dividing total area of all tumor lesions by the total lung area. For the representative comparison between genotypes (*KrasG12V vs KrasG12V;OSKM* mice), the slide with maximum tumor to lung tissue ratio was used for each mouse.

### Statistics

Prism Software (GraphPad, v.9) was used for statistical analysis. All data are displayed as mean ± SD unless otherwise stated. Group allocation was performed in a randomized manner. For normally distributed data with equal variance, statistical significance was determined using two-tailed unpaired Student’s t-test. Welch’s correction was performed for samples with unequal variance. Two-way ANOVA was used to analyze data with two variables, such as tumor growth over time.

A p-value below 0.05 was considered significant and indicated with an asterisk (*p < 0.05, **p < 0.01, ***p < 0.001).

### Data availability

The bulk RNAseq dataset generated to support the findings of this study are available in the Gene Expression Omnibus (GEO) repository, Accession Number XXX (https://www.ncbi.nlm.nih.gov/geo/). All other data will be available from the authors upon request.

## Acknowledgements

We thank Dr Carla P Martins (AstraZeneca, Cambridge, UK) for the provision of L1475luc cells. P.P., V.N.Q. and P.L-F. were supported by predoctoral fellowships from GAIN (IN606A-2017/002, IN606A-2021/016 and ED481A-2020/066, respectively), from Xunta de Galicia. S.D.S.-A. was supported by a postdoctoral fellowship from GAIN (IN606B-2021/011), Xunta de Galicia. Work in the laboratory of M.C. is funded by grants from MCINN/AEI/FEDER, UE (PID2021-125479OB-I00) and GAIN, Xunta de Galicia (IN607D2021/08). DM-E’s laboratory was supported by the CRUK Cambridge Centre Early Detection Program (RG86786), by a CRUK Program Foundation Award (C62187/A29760), by a CRUK Early Detection OHSU Project Award (C62187/A26989), an MRC New Investigator Research Grant (NIRG) (MR/R000530/1), and a Darley/Sands Downing College Fellowship (G109261).

## Author Contributions

JG, MD, JEM, and AD analyzed data; PP, ZZ, DM, VN-Q, VE-S, PL-F, PG, MG, SDS-A, performed experiments and analyzed data; PP, ZZ, MC and DM-E, designed experiments; PP, MC and DM-E, wrote the manuscript; MC and DM-E, supervised the project and secured funding; all the authors read and commented the manuscript.

## Supplementary Figures

**Supplementary Figure 1.**
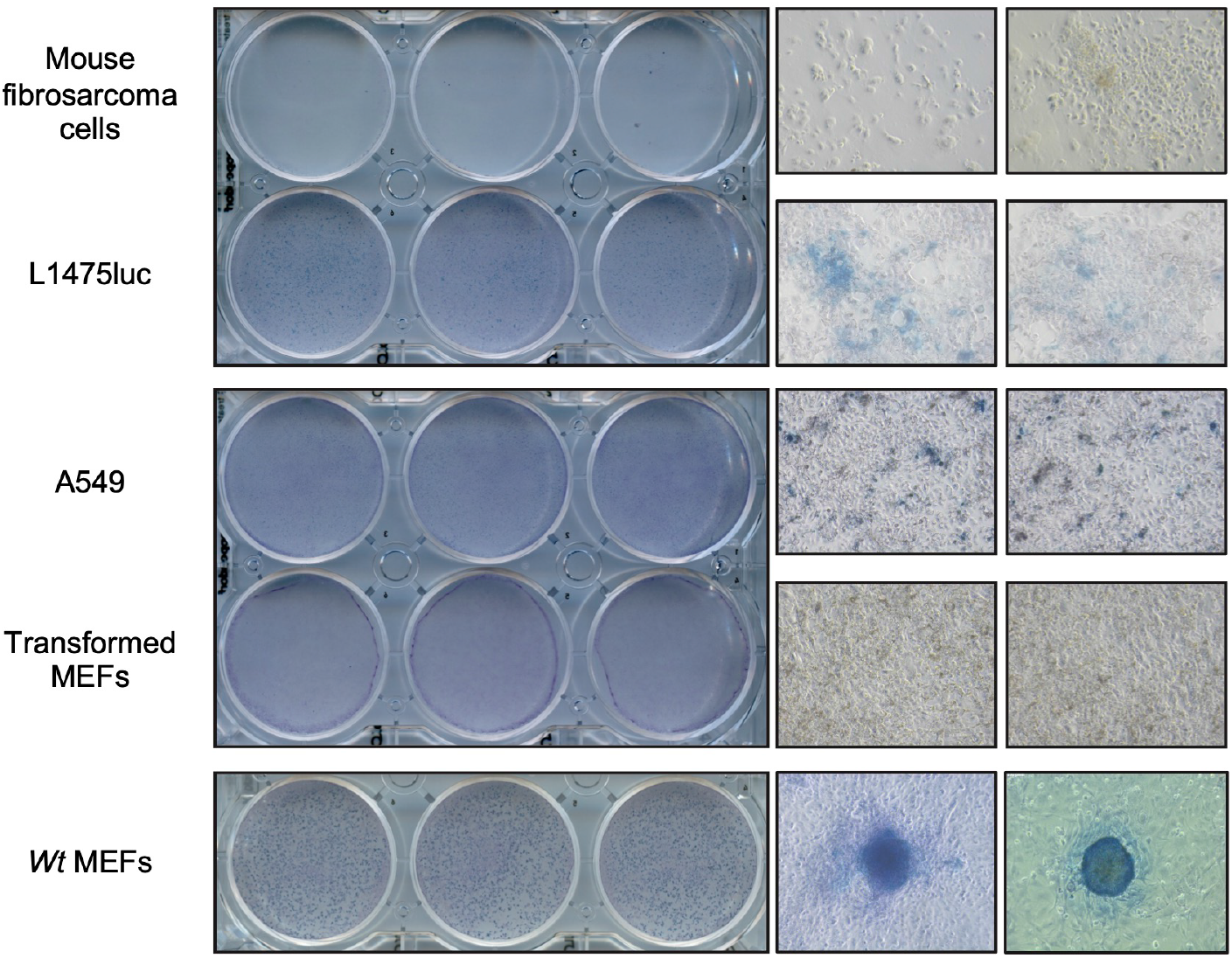
In vitro reprogramming of cancer cell lines and MEFs to iPSCs. **(a)** In vitro reprogramming assays of the indicated cellular types: mouse fibrosarcoma cells, mouse L1475luc and human A549 lung cancer cells, and transformed MEFs (carrying a construct that blocks p53 and another that causes an overexpression of the Ras oncogene). Wild-type (Wt) MEFs were used as positive reprogramming control to iPSCs. Alkaline phosphatase staining plates and representative microscopy images of cells after reprogramming are shown.

**Supplementary Figure 2.**
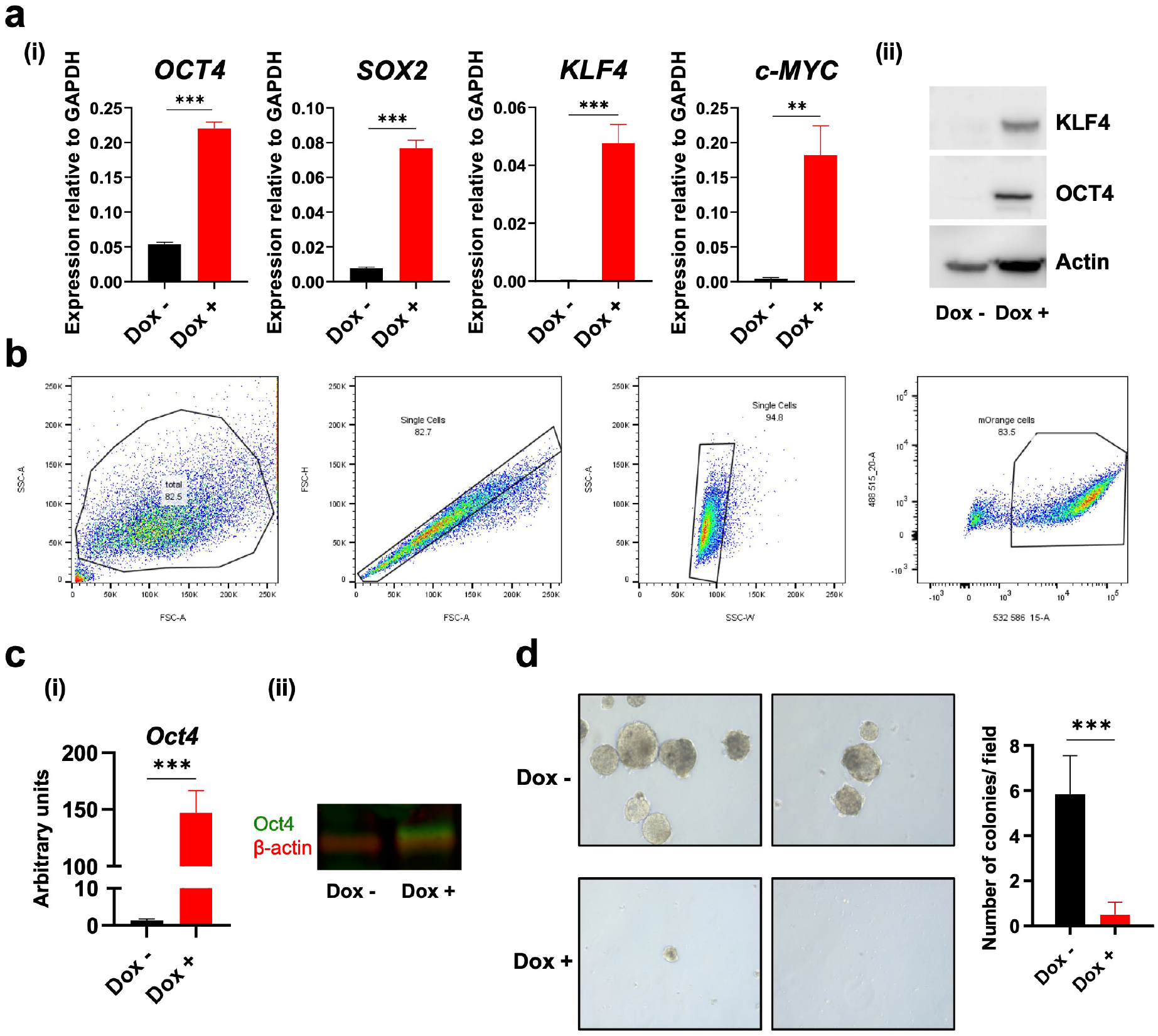
Validation of the expression of reprogramming factors in A549 and L1475luc lung cancer cells. **(a)** mRNA levels of *Oct4*, *Sox2*, *Klf4* and *c-Myc* in A549-rtTA-OSKM cells, treated or not with doxycycline, by RT-qPCR (i), and KLF4 and OCT4 by Western blot (ii). **(b)** Flow cytometry plots showing the gating strategy for the sorting of L1475luc-rtTA-OSKM cells. The gating shows total cells, single cells, alive cells and mOrange positive cells in independent plots. **(c)** Expression levels of Oct4 in L1475luc-rtTA-OSKM cells, treated or not with 1 µg/ml doxycycline, by RT-qPCR (i), and Oct4 by Western blot (ii). Oct4 protein is shown in red color and β-actin in green color. **(d)** Representative image (left) and quantification (right) of a colony formation assay in soft agar of A549-rtTA-OSKM cells treated or not with doxycycline (1µg/ml). Statistical significance was calculated using Student’s t-test, ***P<0.001; **P<0.01; *P<0.05. Data are mean ± SD.

**Supplementary Figure 3.**
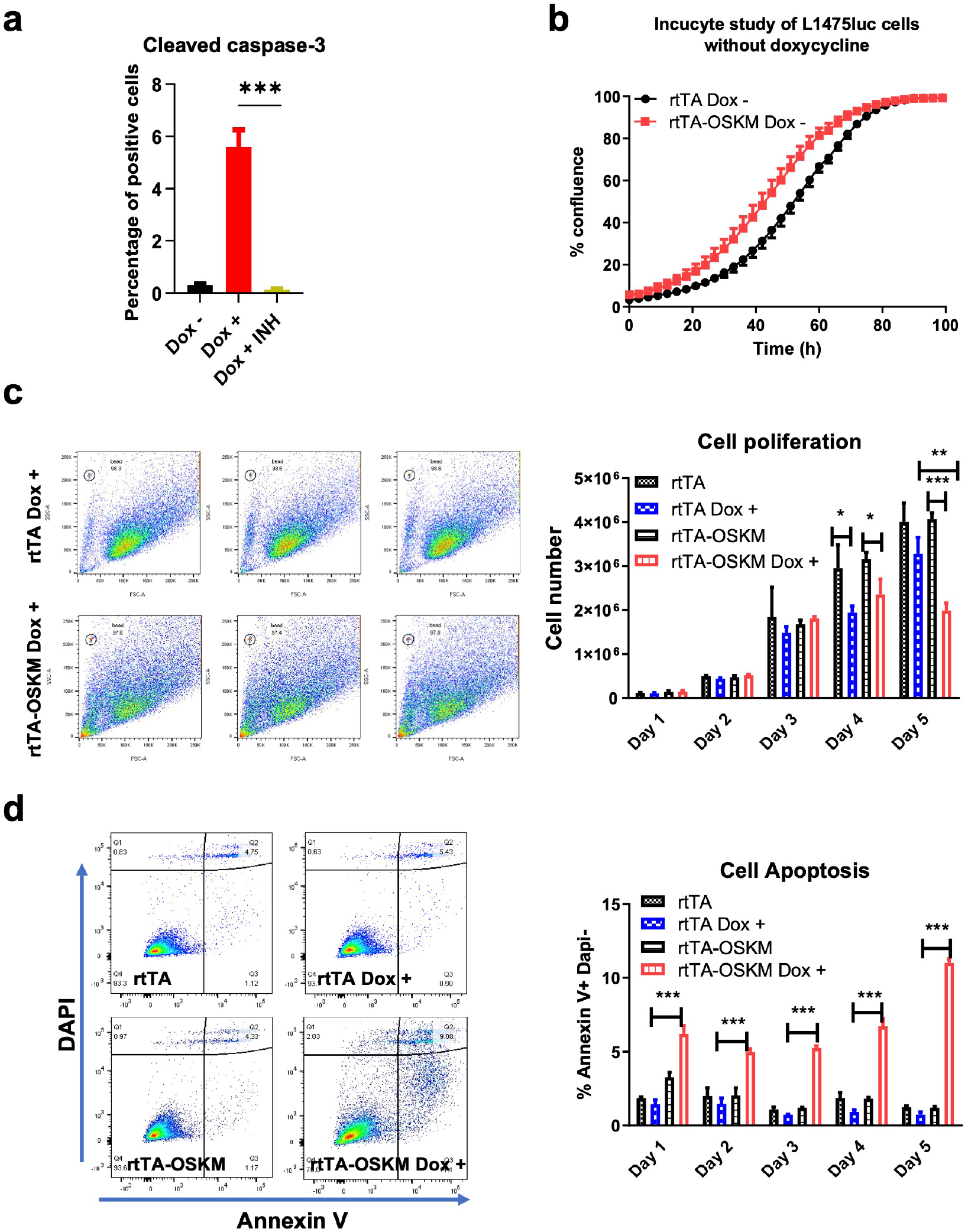
OSKM expression induces lung cancer cell apoptosis and senescence in vitro. **(a)** Percentage of A549-rtTA-OSKM cells positive for cleaved Casapase-3 expressing (Dox +) or not (Dox -) the reprogramming factors, and treated or not with the caspase inhibitor (INH), as indicated. **(b)** Quantification of L1475luc-rtTA and L1475luc-rtTA-OSKM cell proliferation in the absence of doxycycline using Incucyte technology. Images were taken every 3 h for over 96 h. Flow cytometry analysis of L1475luc-rtTA and L1475luc-rtTA-OSKM cell proliferation **(c)** and apoptosis **(d)** in the absence or presence of doxycycline, as indicated. Left, flow cytometry plots. Right, quantification of cell numbers and percentage of Annexin V^+^/DAPI^-^ cells (Q3), as indicated. Statistical significance was calculated using Student’s t-test, ***P<0.001; **P<0.01; *P<0.05. Data are mean ± SD.

**Supplementary Figure 4.**
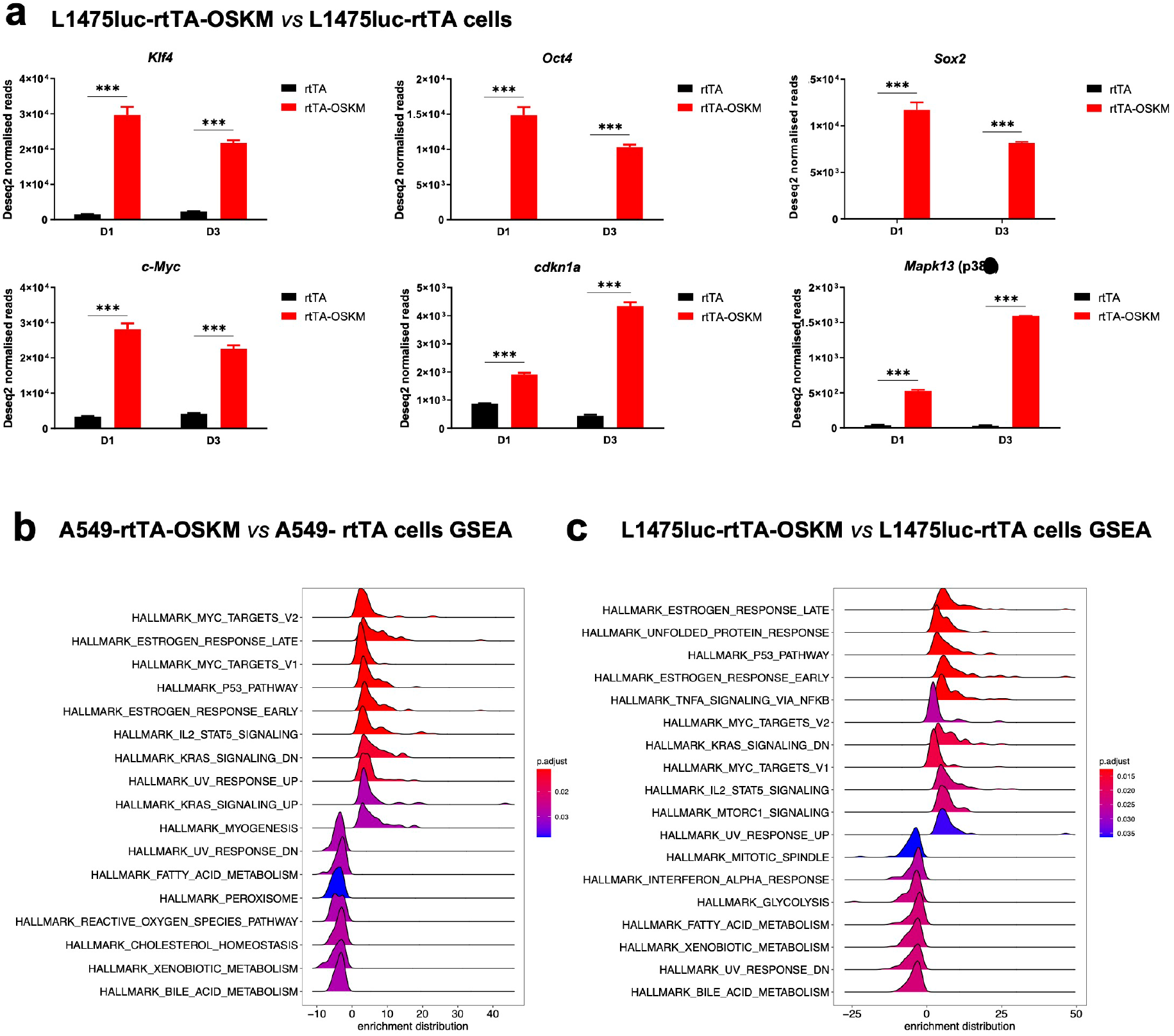
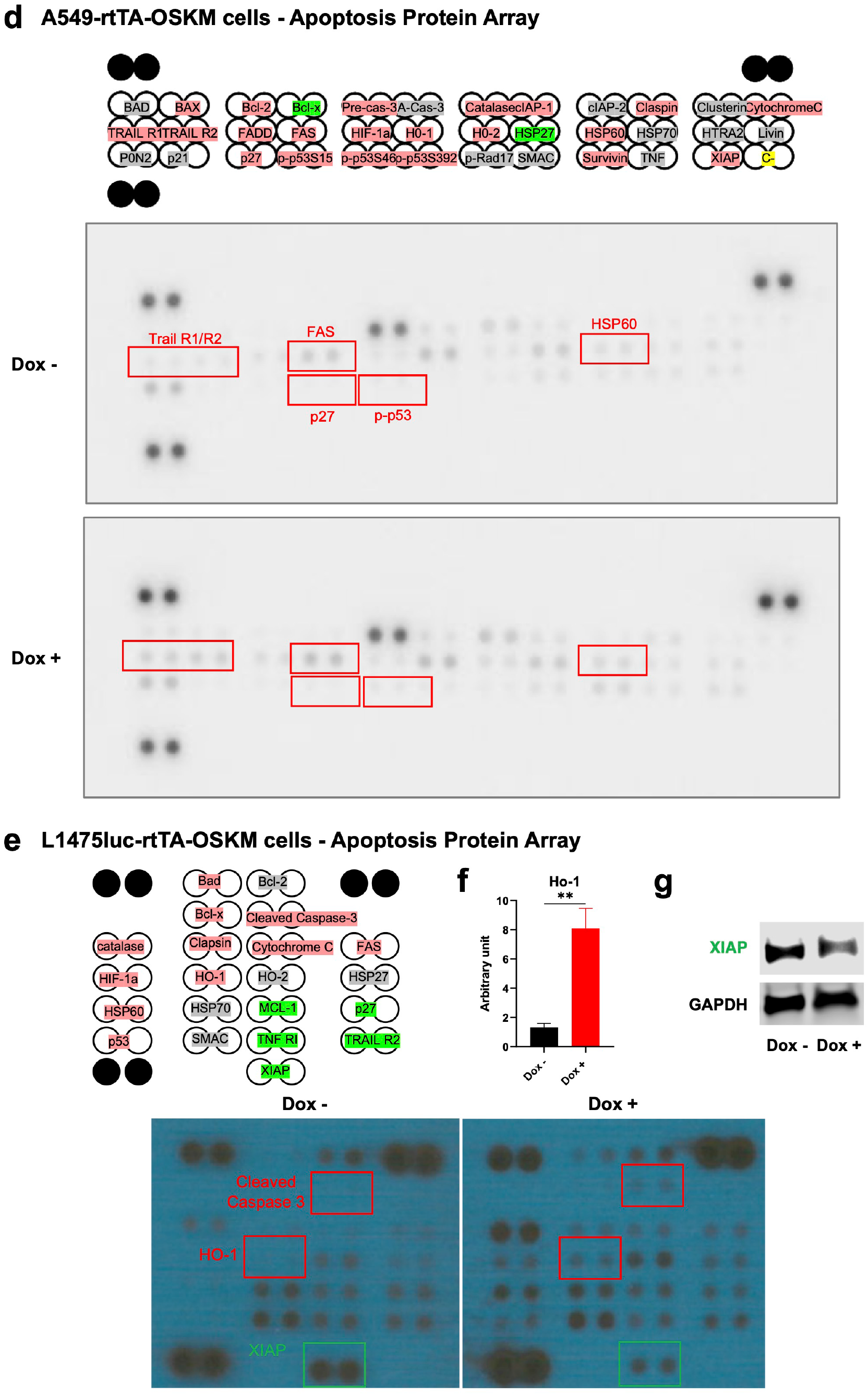
Transcriptomic and proteomic profiles of lung cancer cells upon OSKM expression. **(a)** mRNA levels of *Oct4*, *Sox2*, *Klf4*, *c-Myc*, *Cdkn2a* and *Mapk13* in L1475luc-rtTA and L1475luc-rtTA-OSKM cells at day 1 or 3 post-doxycycline treatment, by RT-qPCR. Ridge plot displaying the gene set enrichment analysis (GSEA) for A549-rtTA-OSKM *vs* A549-rtTA cells **(b)** and L1475luc-rtTA and L1475luc-rtTA-OSKM **(c)** with adjusted *p-value* <0.05. Membrane-based antibody apoptosis array for A549-rtTA-OSKM **(d)** and L1475luc-rtTA-OSKM **(e)** expressing (Dox +) or not (Dox -) the reprogramming factors. Top panel, schematic representation of the array showing capture antibodies sported in duplicate on nitrocellulose membranes for the indicated target apoptotic factors. Red color refers to upregulated protein expression, green color states to downregulated protein expression, and grey color indicates unchanged protein levels. Bottom panels, apoptosis protein array membranes assessing lung cancer cell lines in the presence or absence of doxycycline, as indicated. **(f)** mRNA levels of *Ho-1* in L1475luc-rtTA-OSKM cells treated or not with doxycycline, by RT-qPCR. **(g)** Western blot of XIAP expression levels in L1475luc-rtTA-OSKM cells expressing (Dox +) or not (Dox -) the reprogramming factors. Statistical significance was calculated using Student’s t-test, ***P<0.001; **P<0.01; *P<0.05. Data are mean ± SD.

**Supplementary Figure 5.**
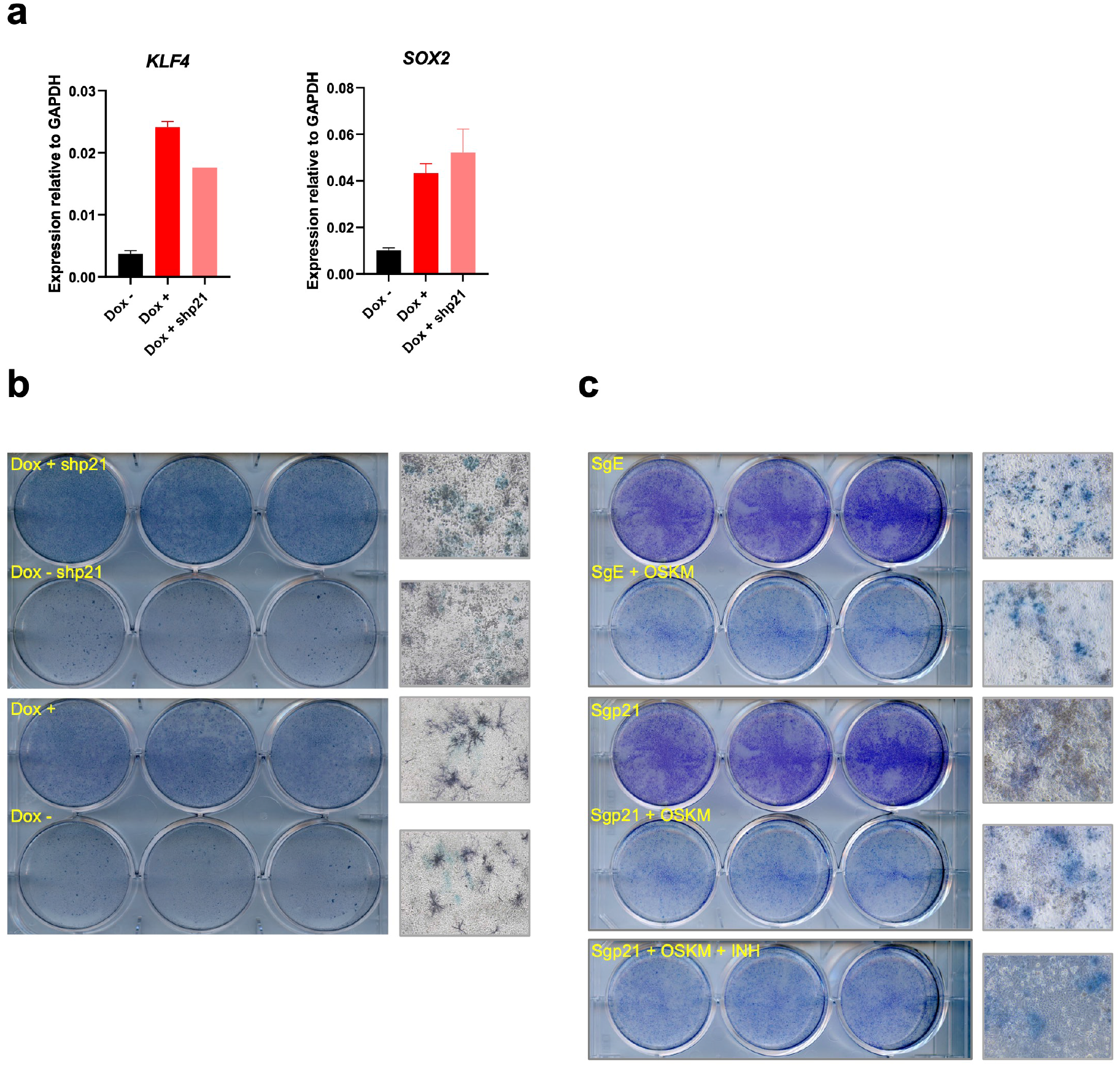
p21 downregulation and knockdown do not allow A549 reprogramming to pluripotency. **(a)** mRNA expression levels of *KLF4* and *SOX2* by RT-qPCR in A549-rtTA-OSKM cells expressing (Dox +) or not (Dox -) the reprogramming factors, and in cells expressing the factors and after knockdown of *p21* (*CDKN1A*) (Dox + shp21). **(b)** In vitro reprogramming assays of A549 cells that do or do not express shp21. Alkaline phosphatase staining plaques and representative microscopy images of cells after reprogramming are shown. **(c)** In vitro reprogramming assays of A549 cells that express CRISPR/Cas9 with an empty guide (SgE), with a guide against p21 (Sgp21), or with a guide against p21 and treated with a pan caspase inhibitor (Sgp21 + INH). Alkaline phosphatase staining plaques and representative microscopy images of cells after reprogramming are shown.

**Supplementary Figure 6.**
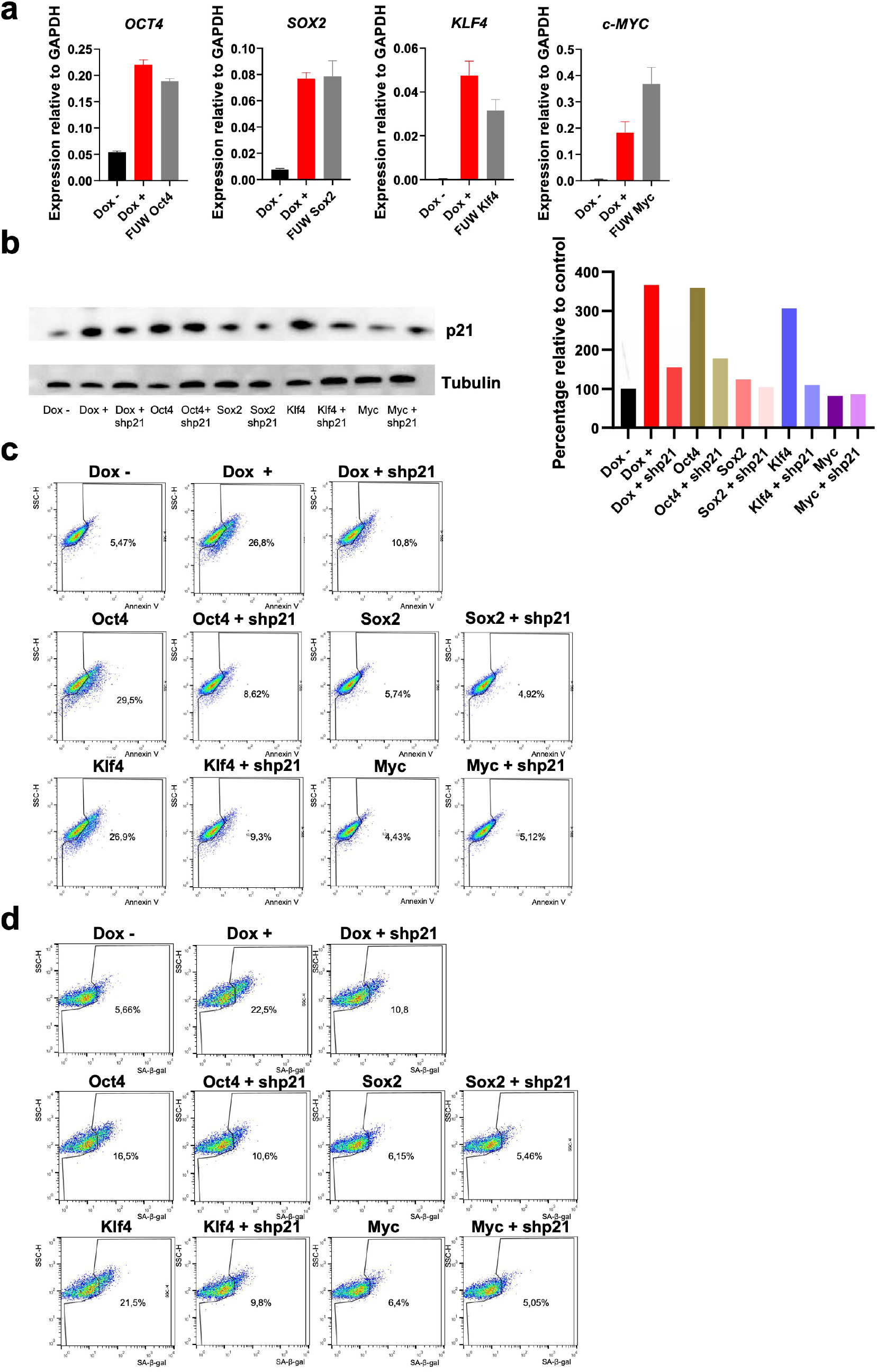
Contribution of individual reprogramming factors to the impaired cell growth and effect of p21 knockdown. **(a)** Expression of reprogramming factors by RT-qPCR in A549 cells that do not express reprogramming factors (Dox -), cells that co-express them (Dox +) and cells that individually express reprogramming factors (FUW OCT4, SOX2, KLF4 or c-MYC). **(b)** Expression of p21 protein levels in A549 cells that express or not the reprogramming factors (Dox +/-), in combination or individually (OCT4, SOX2, KLF4 and c-MYC), alone or after shp21 (+ shp21). Blot (left), quantification (right). **(c)** Flow cytometry plots for Annexin V as a marker indicative of cell apoptosis. **(d)** Flow cytometry plots for SA-β-gal as a marker indicative of cell senescence.

**Supplementary Figure 7.**
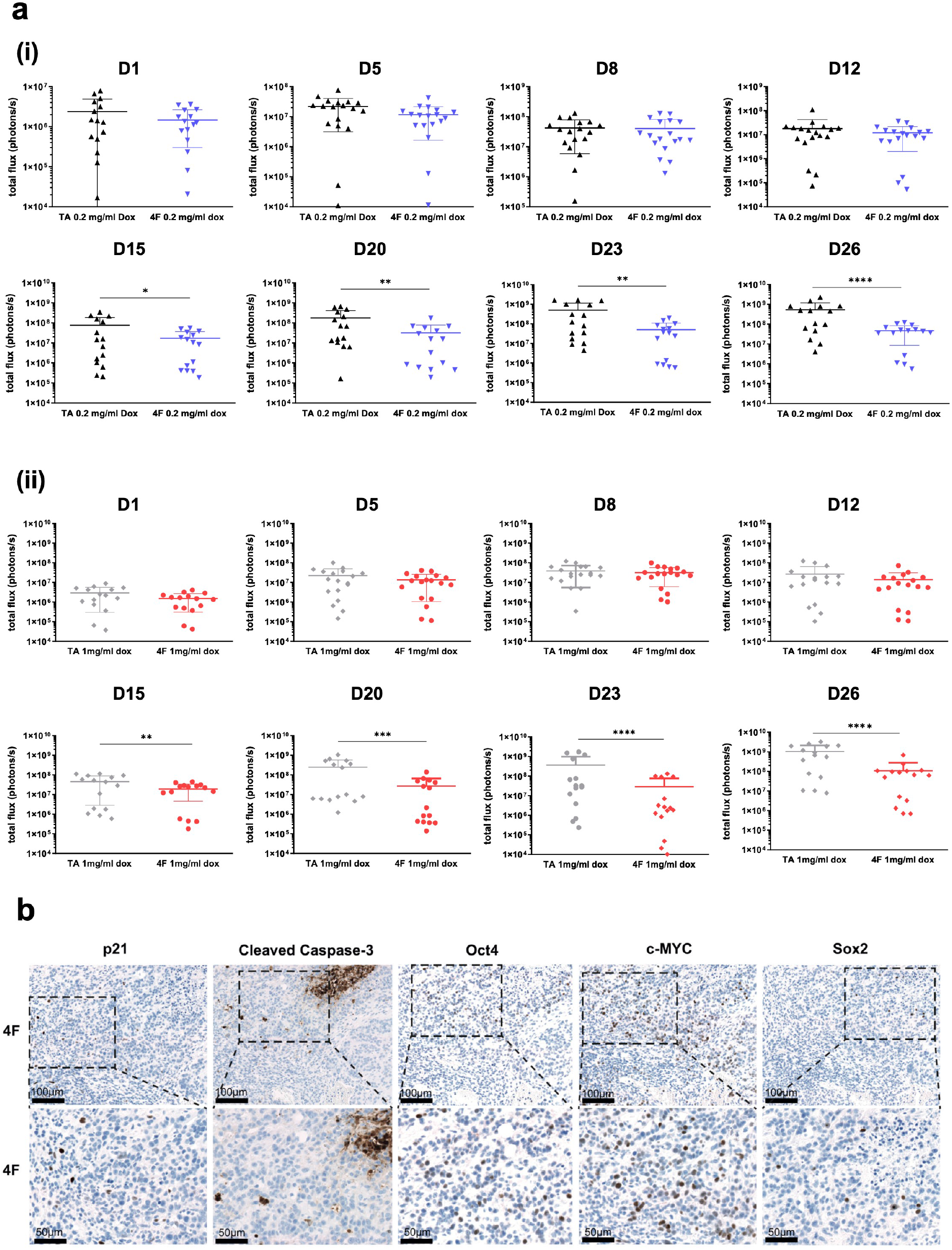
Expression of OSKM impairs growth of subcutaneous tumors. **(a)** Quantification of bioluminescence signal from groups treated with 0.2 mg/ml (i) and 1 mg/ml (ii) doxycycline. **(b)** Immunohistochemistry for p21, cleaved Caspase-3, Oct4, c-Myc and Sox2 in 4F mice. Data shown are mean and SD for each individual day. Wilcoxon matched-pairs signed rank test is used to compare the 2 groups on each day. *, p<0.05; **, p<0.01; ****, p<0.0001.

**Supplementary Figure 8.**
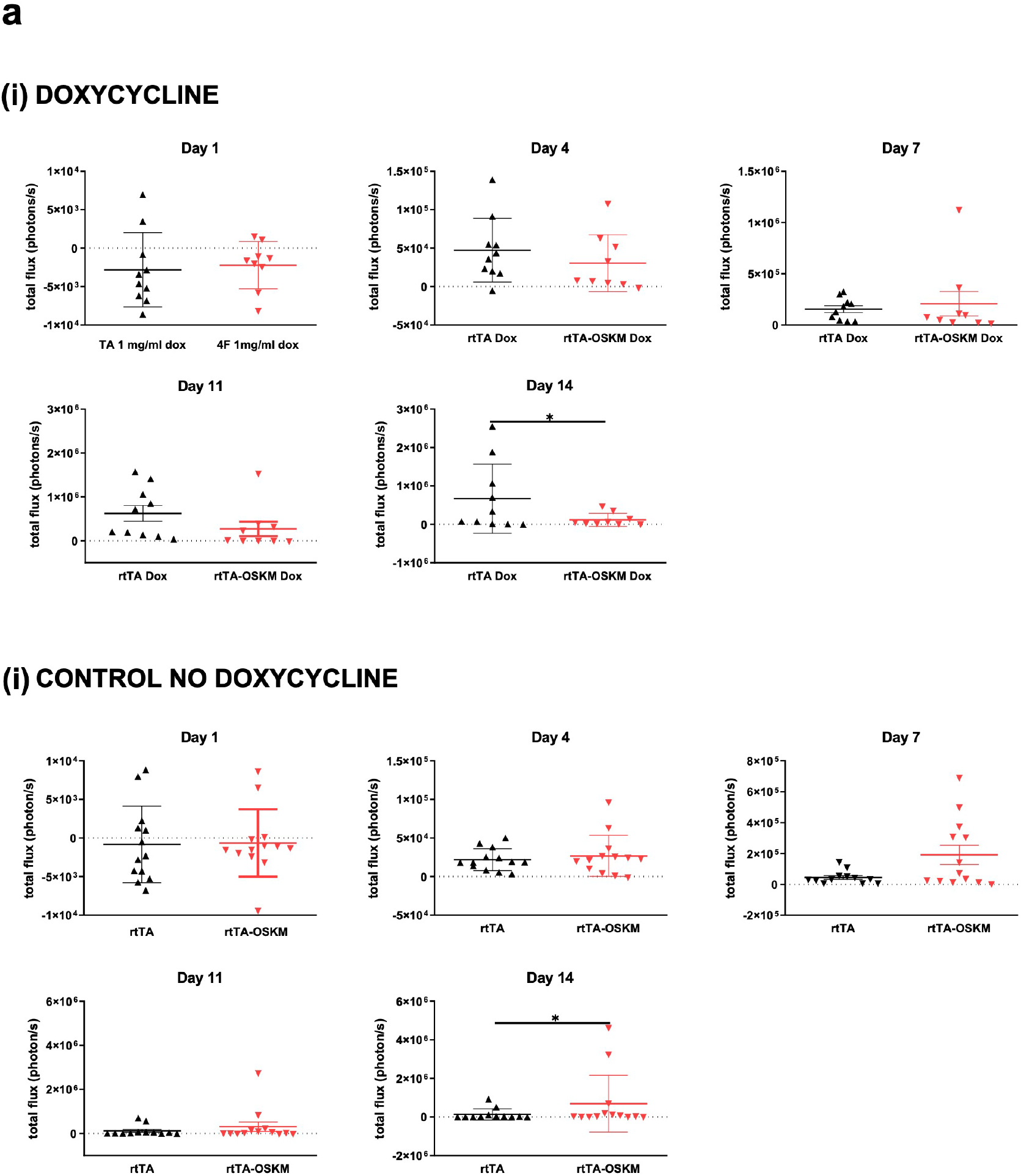
OSKM expression reduces lung tumor burden in a KrasG12V mouse model. **(a)** Quantification of bioluminescence signal from animals treated with 1 mg/ml doxycycline (i) or non-treated (ii). Data shown are mean and SD in each individual day. Wilcoxon matched-pairs signed rank test is used to compare the 2 groups on each day. *, p<0.05; **, p<0.01; ****, p<0.0001.

**Supplementary Figure 9.**
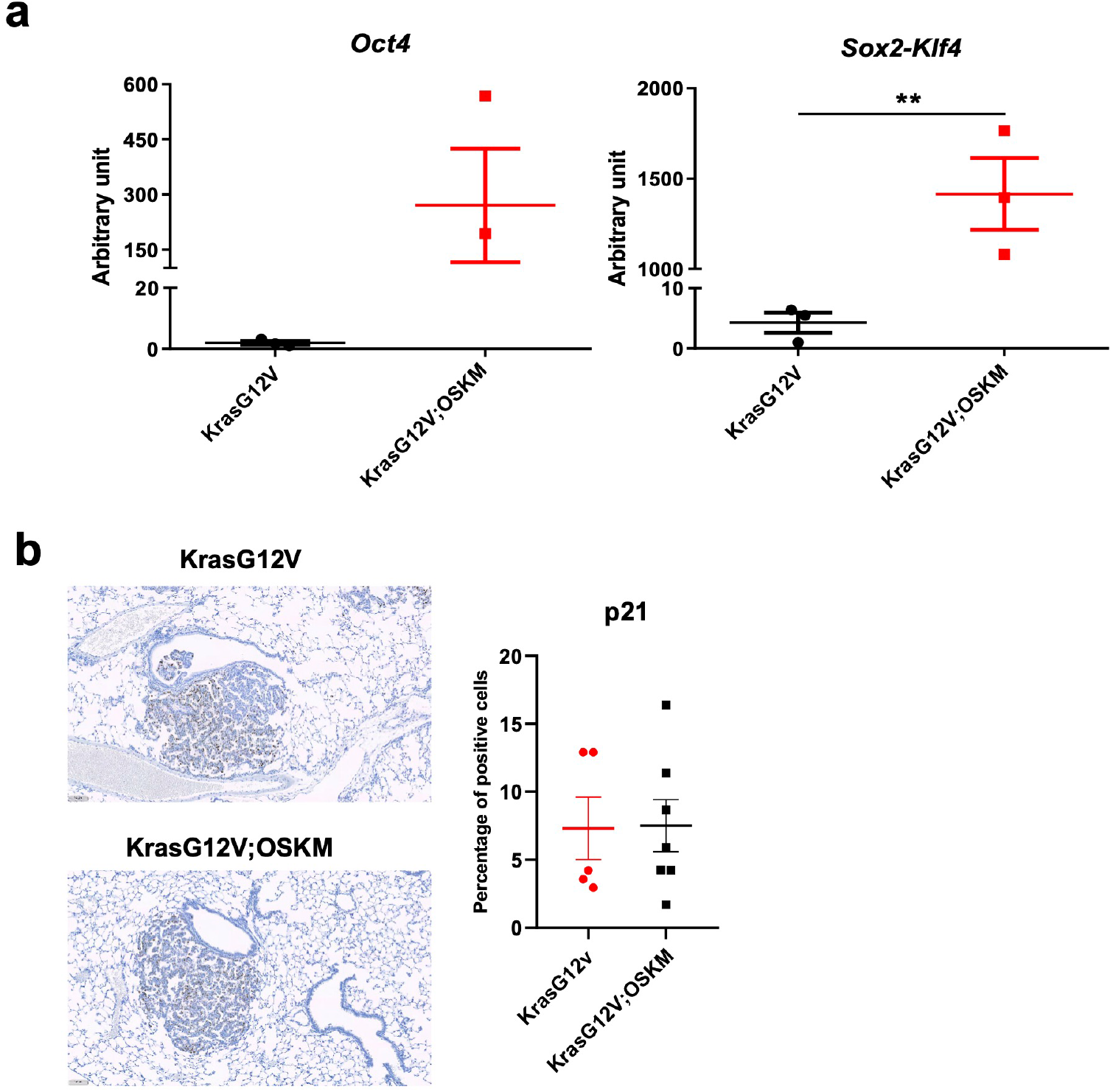
OSKM expression reduces lung tumor burden in a KrasG12V mouse model. **(a)** Oct4 and Sox2-Klf4 expression by RT-qPCR in lungs of mice that express (KrasG12V;OSKM) or not (KrasG12V) the reprogramming factors. **(b)** Immunohistochemistry for p21 in lungs of KrasG12V and KrasG12V-OSKM mice. Representative images (left) and quantification (right).

